# Extracellular Vesicle-Mediated Purinergic Signaling Contributes to Host Microenvironment Plasticity and Metastasis in Triple Negative Breast Cancer

**DOI:** 10.1101/2020.09.11.293837

**Authors:** Suzann Duan, Senny Nordmeier, Aidan E. Byrnes, Iain L. O. Buxton

## Abstract

Metastasis accounts for over 90% of cancer-related deaths. The mechanisms guiding this process remain unclear. Secreted nucleoside diphosphate kinase A and B (NDPK) support breast cancer metastasis. Proteomic evidence confirms their presence in breast cancer-derived extracellular vesicles (EVs). We investigated the role of EV-associated NDPK in modulating the host microenvironment in favor of pre-metastatic niche formation. We measured NDPK expression and activity in EVs isolated from triple-negative breast cancer (MDA-MB-231) and non-tumorigenic mammary epithelial (HME1) cells using flow cytometry, western blot, and ATP assay. We evaluated the effects of EV-associated NDPK on endothelial cell migration, vascular remodeling, and metastasis. We further assessed MDA-MB-231 EV induced-proteomic changes in support of pre-metastatic lung niche formation. NDPK-B expression and phosphotransferase activity were enriched in MDA-MB-231 EVs that promote vascular endothelial cell migration and disrupt monolayer integrity. MDA-MB-231 EV-treated mice demonstrate pulmonary vascular leakage and enhanced experimental lung metastasis, whereas treatment with an NDPK inhibitor or a P2Y1 purinoreceptor antagonist blunts these effects. We identified perturbations to the purinergic signaling pathway in experimental lungs, lending evidence to support a role for EV-associated NDPK-B in lung pre-metastatic niche formation and metastatic outgrowth.

## Introduction

Breast cancer is the second leading cause of cancer death in women, with survival rates heavily influenced by disease stage, tumor grade, and receptor status. While the 5-year relative survival rate for women diagnosed with ductal carcinoma *in situ* remains high, women diagnosed with distant-stage breast cancer see a dramatic decrease in their odds of survival, dropping from a near 99% 5-year survival rate for localized breast cancer to only 27% ^[1]^. This dichotomy in outcomes beckons the need for an improved understanding of the molecular mechanisms that guide breast cancer metastasis and support distant recurrence.

Functional interactions between primary tumor cells and the systemic host microenvironment orchestrate the formation of a metastatic milieu attuned to the establishment and expansion of circulating tumor cells (CTCs). Collectively termed the pre-metastatic niche (PMN), these tissue-specific sites are rich in tumor-supportive factors and demonstrate considerable plasticity and heterogeneity ^[2]^. Cooperative and antagonistic signaling by tumor-secreted factors dictate PMN formation and abet CTCs in immune evasion, extravasation, and distant neocolonization. Such fine-tuned interactions with the distant microenvironment are observed during the early stages of primary tumor development and mediate the escape from indolence to metastatic outgrowth ^[3-5]^. In recent years, tumor derived extracellular vesicles (EVs) have attracted widespread attention for their roles in oncogenesis and are increasingly implicated in multiple cancer hallmarks ^[6,7]^. Further, tumor-derived EVs have been shown to facilitate paracrine-like and organotropic dialogue between primary tumor cells and the PMN ^[8,9]^.

Nucleoside diphosphate kinases (NDPKs) represent a ubiquitous and highly conserved class of multifunctional proteins transcribed from the NME/nm23 gene family and known to be derived from multiple human cancer cell lines ^[10-16]^. Originally identified by Steeg and colleagues as <u>n</u>on-<u>m</u>etastatic gene 23, nm23-H1 (also known as NME1 and NDPK-A) and nm23-H2 (also referred to as NME2 and NDPK-B were among the first genes to be associated with a distinct metastasis suppressor function ^[10,11]^. To date, NDPK-A and NDPK-B have remained the targets of exhaustive research to elucidate their suggested roles as metastasis suppressors. However, they are also associated with multiple cellular functions and may act as nucleoside phosphotransferases, histidine and serine/threonine protein kinases, transcriptional regulators and DNA nucleases ^[12,13]^. Extracellular vesicles derived from multiple human cancer cell lines ^[14-20]^ carry nucleoside diphosphate kinase A and B (EV-associated NDPK-A/B or eNDPK), implicated in promoting angiogenesis and pro-metastatic events extracellularly. The family of NDPK isoforms catalyzes the transfer of a γ-phosphate from a nucleoside triphosphate to a diphospho-nucleoside, and extracellular NDPK localized to the cell surface has been shown to regulate ADP/ATP levels by catalyzing ATP formation from ADP ^[21]^. ATP is a primary agonist to the P2Y and P2X class of purinoreceptors commonly upregulated on tumor cells and vascular endothelium. ATP acting on these purinoreceptors elicits proinflammatory and immunosuppressive responses in the tumor microenvironment and has been shown to promote tumor growth and invasion ^[22-26]^. We have previously demonstrated a pro-angiogenic role for NDPK-mediated activation of P2Y1 receptors on vascular endothelium. Mechanistically, ATP-mediated P2Y1 receptor activation transactivates vascular endothelial growth factor receptor 2 (VEGFR-2) and stimulates mitogenic signaling pathways ^[27-29]^. Additionally, activation of P2Y1/2 receptors on endothelium induces the release of potent vasodilators such as nitric oxide and asserts a mechanistic path for enhanced CTC dissemination ^[30]^. Further, inhibition of NDPK transphosphorylase activity and/or blockade of P2Y1 receptor activation attenuate tumor growth and metastasis to the lung in an orthotopic murine model of triple negative breast cancer ^[31]^.

Extracellular vesicles derived from triple negative breast cancer cells are enriched in NDPK expression, suggesting a functional role for the enzyme in tumor EV-mediated communication with the PMN ^[14-16]^. In support of its purinergic function, elevated ATP in the hundreds micromolar range has been reported in the tumor microenvironment ^[32]^. Moreover, ATP-mediated activation of the P2Y2 receptor was shown to stimulate PMN formation by inducing lysyl oxidase (LOX) and collagen crosslinking ^[33]^. We investigated the hypothesis that NDPK associated with breast cancer EVs propagates purinergic signaling in the distant microenvironment to support PMN formation and metastasis. Here, we show that EVs elaborated by triple negative breast cancer cells are enriched in NDPK-B expression and phosphotransferase activity. We elucidate a functional role for eNDPK-B in the PMN by demonstrating eNDPK-mediated activation of endothelial remodeling in support of angiogenesis and vascular permeabilization. We provide further evidence to support eNDPK-B in metastatic outgrowth and suggest a role for eNDPK-B in modulating the lung vascular microenvironment in favor of PMN formation.

## Materials and Methods

### Drug Reagents

MRS compounds and ellagic acid were purchased from Tocris Bioscience (Bio-Techne, Avonmouth, Bristol, UK). For animal studies, ellagic acid was purchased from MilliporeSigma (Burlington, MA). MRS compounds and ellagic acid were used to antagonize the P2Y1 receptor and inhibit NDPK, respectively.

### Cell Culture of Human Cell Lines

Primary human umbilical vein endothelial cell (HUVEC) and human lung microvascular endothelial cell (HLMVEC) lines (Lifeline Cell Technology, Frederick, MD) were grown in VascuLife-VEGF and Vasculife-MV media, respectively (Lifeline Cell Technology). Media was supplemented with their respective growth factor kits and 10% EV-depleted FBS (Atlanta Biologicals, Flowery Branch, GA). MDA-MB-231, MDA-MB-231-Luc+, MDA-MB-157, MDA-MB-468, MCF-7, MCF-10A, MCF-12F, telomerized human mammary epithelial cells (HME1/HMECs), and 3T3 fibroblast cells were purchased from ATCC (Manassas, VA). MDA-MB-231 and MDA-MB-231-Luc+ cells were transduced with lentivirus vector expressing a GFP or RFP tag fused to the membrane-associated tetraspanin CD63 (System Biosciences, Palo Alto, CA). Breast cancer cells were cultured in Dulbecco’s Modified Eagle Medium (DMEM) supplemented with 2% penicillin-streptomycin and 10% EV-depleted FBS. FBS was filtered through a 0.2 μm membrane and ultracentrifuged at 100,000 × *g* for 24 hours at 4°C to deplete EVs. Non-transformed breast epithelial cell lines were cultured in Human Mammary Epithelial Cell Medium (HuMEC) supplemented with 2% penicillin-streptomycin, 10% EV-depleted FBS, and HuMEC supplement kit excluding bovine pituitary extract (Thermo Fisher Scientific, Waltham, MA). Cells were cultured under 5% CO2 in room air at 37°C.

### Neonatal pulmonary endothelial cell isolation

Approval was granted by the Institutional Animal Care and Use Committee prior to animal use. Methods were adapted from Sobczak *et al* ^[40]^. Digested lung tissue was incubated with Dynabeads™ (Thermo Fisher Scientific) coated with intercellular adhesion molecule (ICAM-1, ICAM-2) and vascular cell adhesion molecule (VCAM-1) antibodies conjugated to Alexa Fluor 680 (BioLegend, San Diego, CA). Sorted cells were cultured in Endothelial Growth Media (EGM-2) (Lonza, Basel, Switzerland) on 2% gelatin-coated chamber slides. Cells were fixed in 4% paraformaldehyde (PFA), incubated in anti-wheat germ agglutinin conjugated to Alex Fluor 488 (1:1000, Thermo Fisher Scientific), and imaged by CLSM at 200X magnification.

### Extracellular Vesicle Isolation

Cells were cultured for 4-5 days in DMEM with 10% EV-depleted FBS. Conditioned media was centrifuged at 3,000 × *g* for 10 minutes and the supernatant was filtered through a 0.2 μm membrane before centrifuging again at 20,000 × *g* for 30 minutes. The supernatant was ultracentrifuged at 100,000 × *g* for 70 minutes and the EV pellet was washed in PBS before repeating the spin. EVs were re-suspended in sterile PBS and total protein was quantified using an EZQ assay (Thermo Fisher Scientific). For data presented in supplemental figures 1 and 2, EVs were isolated using ExoQuick-TC (System Biosciences) according to manufacturers’ instructions. For experimental metastasis study only, both ultracentrifugation- and ExoQuick-TC-purified EVs were used in order to meet experimental demand (four weeks treatment with each).

### Transmission Electron Microscopy (TEM)

Transmission electron microscopy grids were prepared according to Thery *et al* ^[35]^. Grids were analyzed by the University of Nevada, Reno Imaging Core using the Philips CM10 transmission electron microscope and a GATAN BioScan 792 digital scanner.

### EV Analysis with Flow Cytometry

FACS protocol by Thery *et al* ^[35]^ was used for flow cytometry analysis of EVs adsorbed to latex beads. Briefly, purified EVs were incubated with 3.9 μm latex beads (Thermo Fisher Scientific) and rocked overnight at 4°C. Beads with adsorbed EV proteins were blocked in 100 mM glycine for 30 min at room temperature, then centrifuged for 3 min at 1500 x g. Bead pellet was washed three times with 0.5% BSA in PBS before incubating in 1:200 rabbit anti-human NM23A (Abcam), NME2 (Abcam), or matched rabbit isotype primary antibodies (Abcam) for 60 min at 4°C. Beads were washed twice and incubated in PE-conjugated donkey anti-rabbit secondary antibody (Biolegend) for 30 min at 4°C. Samples were washed twice, resuspended in 0.5% BSA and analyzed on the BD LSR II Flow Cytometer (BD Biosciences, San Jose, CA) at the University of Nevada, Reno Flow Cytometry Core. For surface labeling, EVs were incubated for 30 min at room temperature with PE-conjugated mouse IgG1 isotype (Biolegend), CD63 (Biolegend), or NDPK-A (US Biologicals) antibodies diluted in 0.5% BSA in PBS. Data was analyzed using FlowJo software version 10.5.3. For NDPK-B quantitation, side scatter area (SSC-A, y-axis) was plotted against PE area (PE-A, x-axis) on a log scale. Gates were manually drawn to exclude 98-99% of sample isotype signal intensity and copied to respective plots of primary antibody labeling. Gates captured approximately 95% of total NDPK-B signal. Overlapping values from respective isotype controls were subtracted from gated NDPK-B and CD63 values and NDPK-B was normalized to CD63 intensity and plotted using Graphpad Prism version 8 (n = 2 independent experiments).

### Western Blot Analysis

For NDPK expression analysis, EV protein was mixed with SDS sample buffer and 1% triton-X before heating at 70°C for 10 min. Samples were loaded onto a 4-20% pre-cast TGX polyacrylamide gel (BioRad) and run at 200V for approximately 45 min. Proteins were transferred onto a 0.2 μm nitrocellulose membrane using the Turboblot transfer system. Membranes were blocked in 5% BSA in PBS with 0.05% Tween-20 and incubated overnight at 4°C in NM23A (Abcam), NME2 (Abcam), and CD9 (Biolegend) primary antibodies diluted 1:250. Membranes were imaged on the Licor Odyssey with Image Studio software. For 10 μg of EV protein and cell lysate were prepared as previously described. HUVEC and lung lysates were prepared as described previously, except in reducing conditions. Antibody dilutions and sources are listed in Supplementary Materials.

### Wes Protein Assay

HUVECs were treated for three hours with 100 ng/ml HME1 or MDA-MB-231 EVs, alone or with 100 uM MRS2179 and ellagic acid. Equal protein lysate concentrations (0.2 mg/ml) were run on ProteinSimple Wes™ according to manufacturer instructions (Biotechne, Minneapolis, MN). eNOS (1:100, Santa Cruz Biotechnology, Dallas, TX), iNOS (1:50, Santa Cruz Biotechnology), COX-2 (1:50, Cell Signaling Technologies), and GAPDH (Cell Signaling Technologies) antibodies were used to measure protein expression.

### NDPK Transphosphorylation Activity Assay

EV lysates were prepared in 10% triton X-100 in room Air Krebs (RAK) buffer without CaCl2. Recombinant NDPK-A, NDPK-B, and ATP were serially diluted into standards and incubated with 10 µM ADP and 30 µM UTP before quenching with 0.1 M HCl in prepared buffer. Standards were run with whole EVs or EV lysates. The pH was neutralized to 7.4 and equal volumes of luciferin-luciferase ATP detection buffer were added. Luminescence was measured using the Hidex Chameleon plate reader with MikroWin 2000 software.

### Transwell Migration Assay

Transwell inserts with 3 μm pores and 12 mm diameter (Corning, Corning, NY) were coated with rat-tail collagen (Corning). HUVECs were grown to 75% confluence and serum-starved for 24 hours prior to seeding in top chambers containing VascuLife media with 2% EV-depleted FBS (2.5 × 10^5^ cells per well). Assay was initiated by adding EVs (100 μg/ml or 1 ug/ml) to bottom chambers and P2Y1 receptor antagonists (10 μM MRS2179, 100 nM MRS2279, 10 nM MRS2500) or an NDPK inhibitor (10 μM EA) to top chambers (n = 3). After 24 hours, membranes were processed and imaged at 100X magnification using the Keyence BZ-X700 automated microscope. Migrated cells were quantified using the BZ-X700 Analyzer Software (Keyence, Osaka, Japan). Three images at the center of each well were quantified and averaged to generate a count for each well. Assay was repeated with MLECs in EGM-2 media (Lonza).

### Transwell Permeability Assay

Methods were adapted from Martins-Green *et al* ^[32]^. Transwell inserts were coated with growth factor-reduced Matrigel® (Corning) and seeded with HUVECs and HLMVECs in Vasculife-VEGF or Vasculife-MV media (1 × 10^5^ cells/well). A second monolayer was seeded 48 hours later. Assay was initiated 48 hours later with the addition of 10 μg/ml final concentration of FITC-dextran sulfate (MilliporeSigma) and 100 ng/ml EVs to lower chambers, while inhibitors (100 μM EA, MRS2179, or both) were added to top chambers. Media was collected from top chambers at indicated time points and fluorescence was measured with 485 nm excitation/535 nm emission using the Hidex Chameleon plate reader and MikroWin 2000 software. n = 6.

### Imaging of Endothelial Barrier

HLMVECs were seeded and treated as described in the previous experiment, but without the addition of FITC-dextran. Cells were treated for three hours, washed with PBS, fixed in 4% PFA, permeabilized with 1% Triton-X, and incubated overnight in 1:250 FITC-phalloidin (Thermo Fisher Scientific) or ZO-1 and β-catenin antibodies (Thermo Fisher Scientific). Whole membranes were mounted with Vectashield containing DAPI (Vector Laboratories, Burlingame, CA). Slides were imaged by CLSM at 200X magnification.

### Confocal Microscopy

Image acquisition and analysis were performed using an Olympus IX81 Fluoview confocal microscope and FV10-ASW software (version 4.02, Windows 7, Olympus America, Inc., Melville, NY). All image acquisition settings and post hoc image adjustments (i.e. brightness, contrast, and LUT) were applied globally for each experiment to ensure accuracy in comparison between controls and experimental groups.

### Animal Studies

The University of Nevada, Reno Institutional Animal Care and Use Committee approved all studies and procedures prior to animal use. NCr/SCID mice were purchased from Charles River Laboratories (Wilmington, MA) and bred on site. All subsequent litters were age matched for experiments.

### In Vivo Vesicle Tracking

7-week old female SCID mice (n = 2) were orthotopically injected in the mammary fat pad with CD63-GFP-labeled MDA-MB-231-Luc^+^ cells (2 × 10^5^ cells) suspended in DMEM and Matrigel®. Mice were imaged bi-weekly using Living Image Software Version 4.5 and the Lumina III *In Vivo* Imaging System (IVIS, PerkinElmer, Waltham, MA). Serum was collected every other week and imaged by CLSM at 200X magnification to detect circulating GFP-labeled EVs. Lungs were collected at nine weeks, embedded, and cryo-sectioned to a 10 µm thickness. Slides were fixed with 4% PFA, stained with wheat germ agglutinin conjugated to 1:100 Texas Red, and imaged by CLSM at 400X magnification. EVs were detected using a 1:100 anti-GFP Tag antibody (Thermo Fisher Scientific, #GF28R).

### TMT-labeled Mass Spectrometry

8-week old female SCID mice were injected by tail vein with 5-10 μg of purified MDA-MB-231 EVs three times per week for three weeks. Whole lungs were collected, powdered, and reconstituted in NP-40 lysis buffer. Samples were submitted to the Nevada Proteomics Center for mass spectrometry analysis. Methodology and analysis is described in detail in Supplementary Materials.

### Evans Blue Dye (EBD) Extravasation Study

10-week old SCID mice were injected by tail vein with PBS vehicle or 5-10 μg of purified MDA-MB-231 EVs three times per week for eight weeks. MRS2179 was injected by tail vein with EVs (8.5 μg) while ellagic acid (EA) was administered via drinking water (120 μg/ml equivalent to 5 ± 0.7 ml/day per mouse ^[29]^). A final group received both drugs (Combo). After 8 weeks, mice were injected by tail vein with 2% EBD and re-caged for three hours (n = 4-6 per group). Lungs were collected following cardiac perfusion with PBS. Right lung lobes were incubated in *N,N*-dimethylformamide to extract EBD. Absorbance at 610 nm was measured using MikroWin 2000 software. Left lung lobes were cryo-sectioned to 10 μm thickness and imaged by CLSM at 100X magnification.

### Experimental Metastasis Study

10-week old SCID mice were treated for eight weeks as previously described (n = 3-5 per group). Following treatment, mice were injected by tail vein with 1×10^5^ MDA-MB-231-luc^+^ cells engineered to express GFP-tagged CD63. Mice were imaged weekly using the Lumina III *In Vivo* Imaging System (IVIS, PerkinElmer). Thirty days later, whole lungs, liver, and brain were removed, incubated in D-luciferin, and imaged.

### Statistical Analysis

One- or two-way analysis of variance (ANOVA) was used for assays comparing three or more groups. Unpaired two-tailed Student’s t-test or Mann-Whitney nonparametric test were used for all other assays comparing between two groups, as indicated in figure legends. Tukey posttest was applied when appropriate, and significance was defined as follows: * = *p* < 0.05, ** = *p* < 0.01, *** = *p* <0.001.

## Results

### Extracellular vesicles secreted by non-tumorigenic human mammary cells and triple negative breast cancer cells demonstrate classical morphological and molecular features

EVs were purified by ultracentrifugation from the conditioned media of triple negative human breast cancer cells (MDA-MB-231) and from a non-tumorigenic mammary epithelial cell line (HME1). TEM imaging revealed extracellular vesicles that were predominantly 50 to 80 nm in diameter. EVs possessed a characteristic phospholipid bilayer membrane and were relatively uniform in size and morphology (Figure 1A). Compared to cell lysates, EVs were enriched in classical exosome and EV markers, including CD9, CD63, CD81, ALIX, flotillin-1, and MFGE8. In contrast, cell lysates were enriched in calnexin, a cellular protein known to be absent in pure EV preparations (Figure 1B). EVs from MDA-MB-231 cells exhibited lower expression of the EV marker tsg101 compared to cell lysates, while HME1-derived EVs expressed low levels of calnexin. For downstream visualization and tracking studies, MDA-MB-231 cells were transduced with lentivirus expressing GFP- or RFP-labeled CD63. Expression of GFP and RFP EVs was confirmed by confocal imaging (Figure 1C). EVs were also purified using ExoQuick-TC and further characterized by TEM and flow cytometry. EVs that were isolated using ExoQuick-TC were approximately 30-100 nm in diameter, displayed a phospholipid bilayer, and demonstrated positive labeling for CD63, CD9, and CD81 (Supplemental Fig. 1).

**Figure 1.**
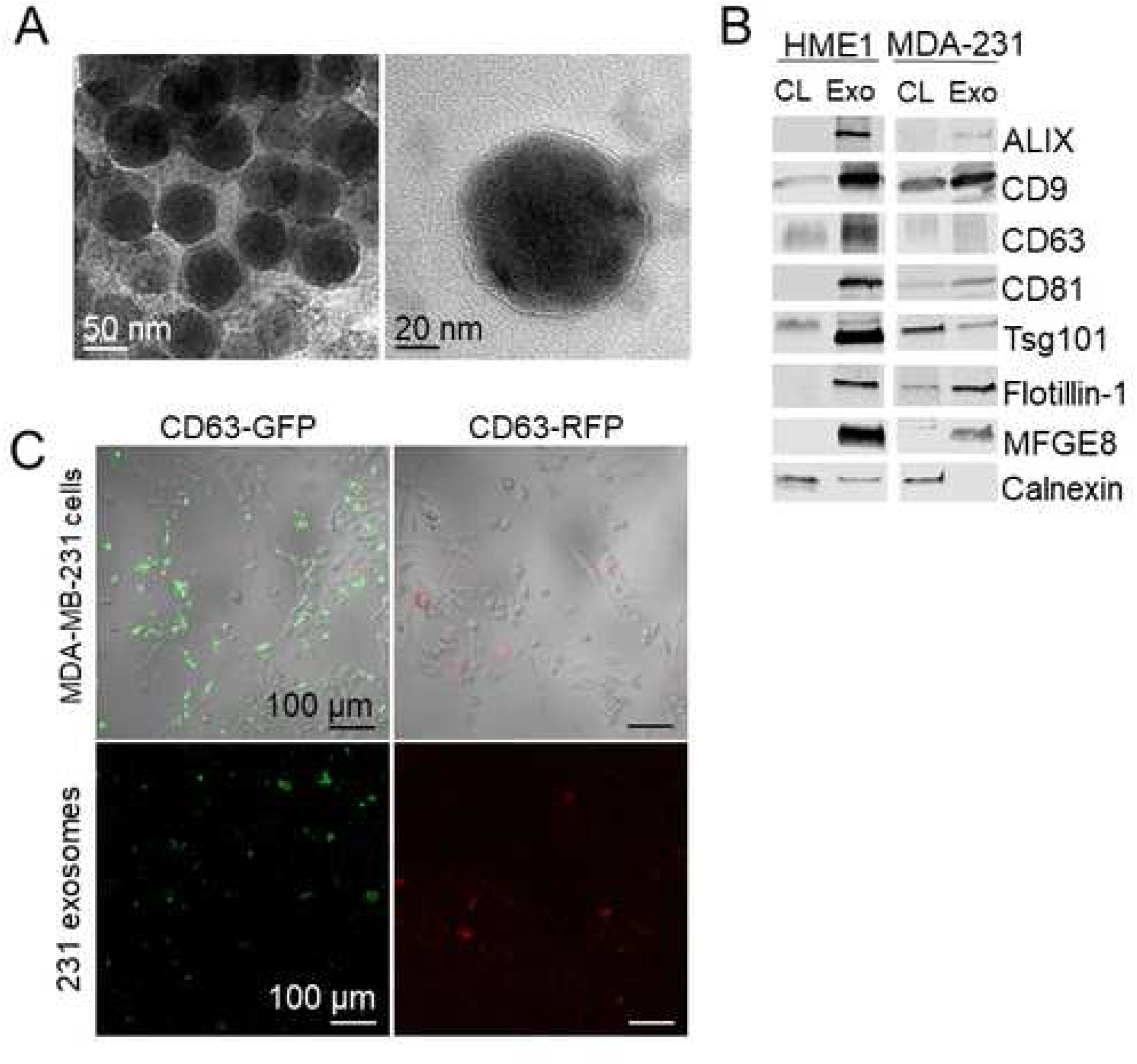
Characterization of HME1 and MDA-MB-231 EVs by transmission electron microscopy, confocal microscopy, and western blot. *(A)* TEM images of MDA-MB-231 EVs stained with uranyl-acetate. Scale 50 and 20 nm. *(B)* Western blots of classical exosome and EV protein markers in HME1 and MDA-MB-231 EV lysates. Calnexin was evaluated as a cell-enriched marker. *(C)* CLSM images of MDA-MB-231 cells and respective purified EVs expressing GFP or RFP-labeled CD63. Scale 100 μm.

### MDA-MB-231 EVs are enriched in NDPK-B expression compared to non-tumorigenic mammary EVs

To evaluate whether HME1 and MDA-MB-231 EVs carry NDPK-A and NDPK-B, EVs were adsorbed to latex beads and analyzed by flow cytometry. HME1 and MDA-MB-231 EVs demonstrated positive labeling for CD63. Consistent with the previous western blot analysis, HME1-derived EV preparations exhibited higher CD63 expression compared to EV preparations from MDA-MB-231 cells (Figure 2A). Both populations demonstrated robust labeling for NDPK-B and minimal labeling for NDPK-A compared to isotype control (Figure 2B). Comparatively low surface expression of CD63 and NDPK-A/B on non-adsorbed EVs was further confirmed by positive antibody labeling (Figure 2C). As CD63 is the most frequently used marker to define bona fide EV populations, we used the expression of the tetraspanin as an internal quantitative control for total EV protein. Relative to CD63, NDPK-B expression was higher in MDA-MB-231 EVs compared to HME1 EVs (Figure 2D). Consistent with these observations, western blot analysis confirmed more robust CD9 expression in HME1 EVs compared to MDA-MB-231 EVs. Relative to CD9, NDPK-A and NDPK-B expression was approximately three-fold higher in MDA-MB-231 EVs. Moreover, EVs derived from both cell lines possessed nearly 30-fold higher expression of NDPK-B compared to NDPK-A (Figure 2E)

**Figure 2.**
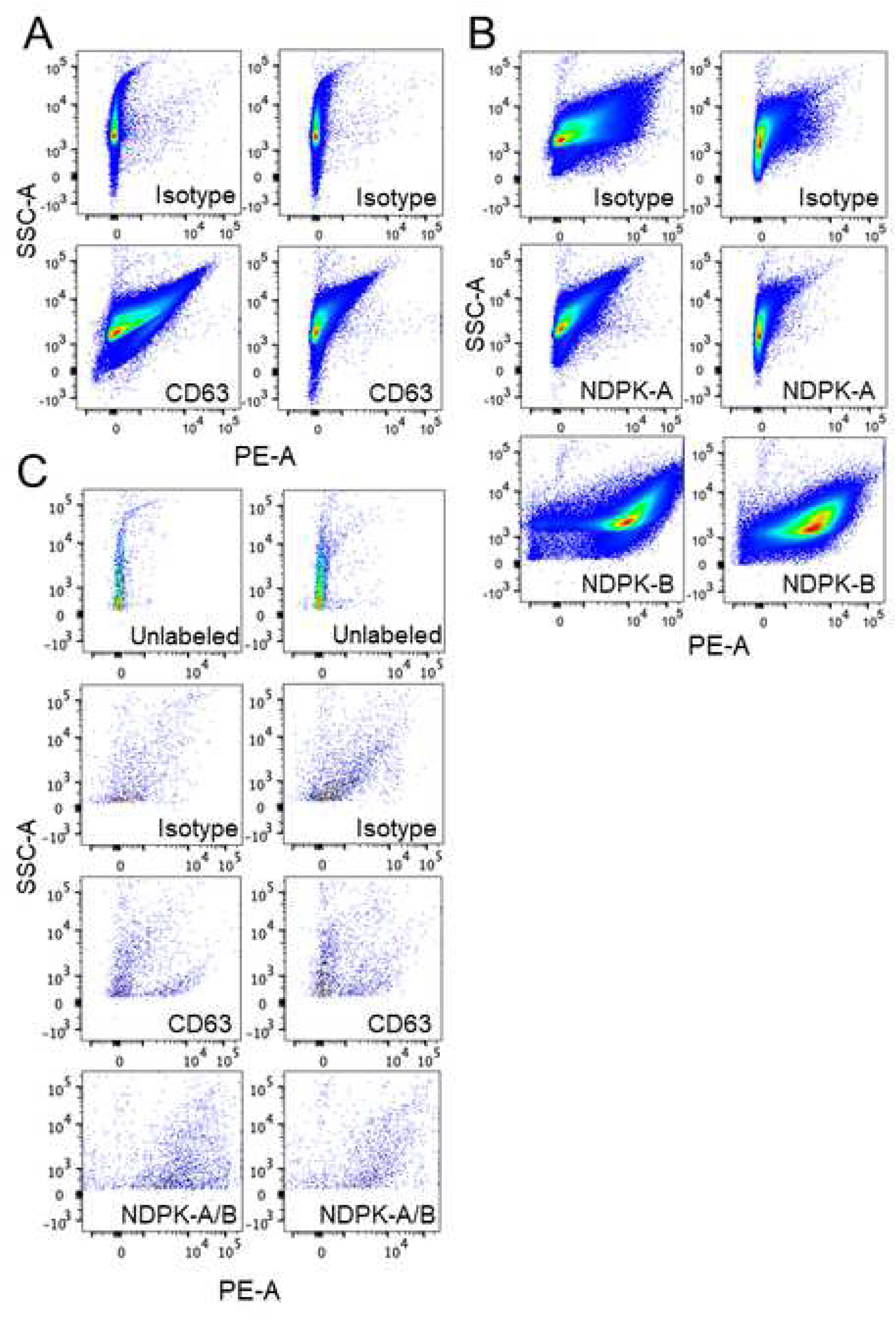
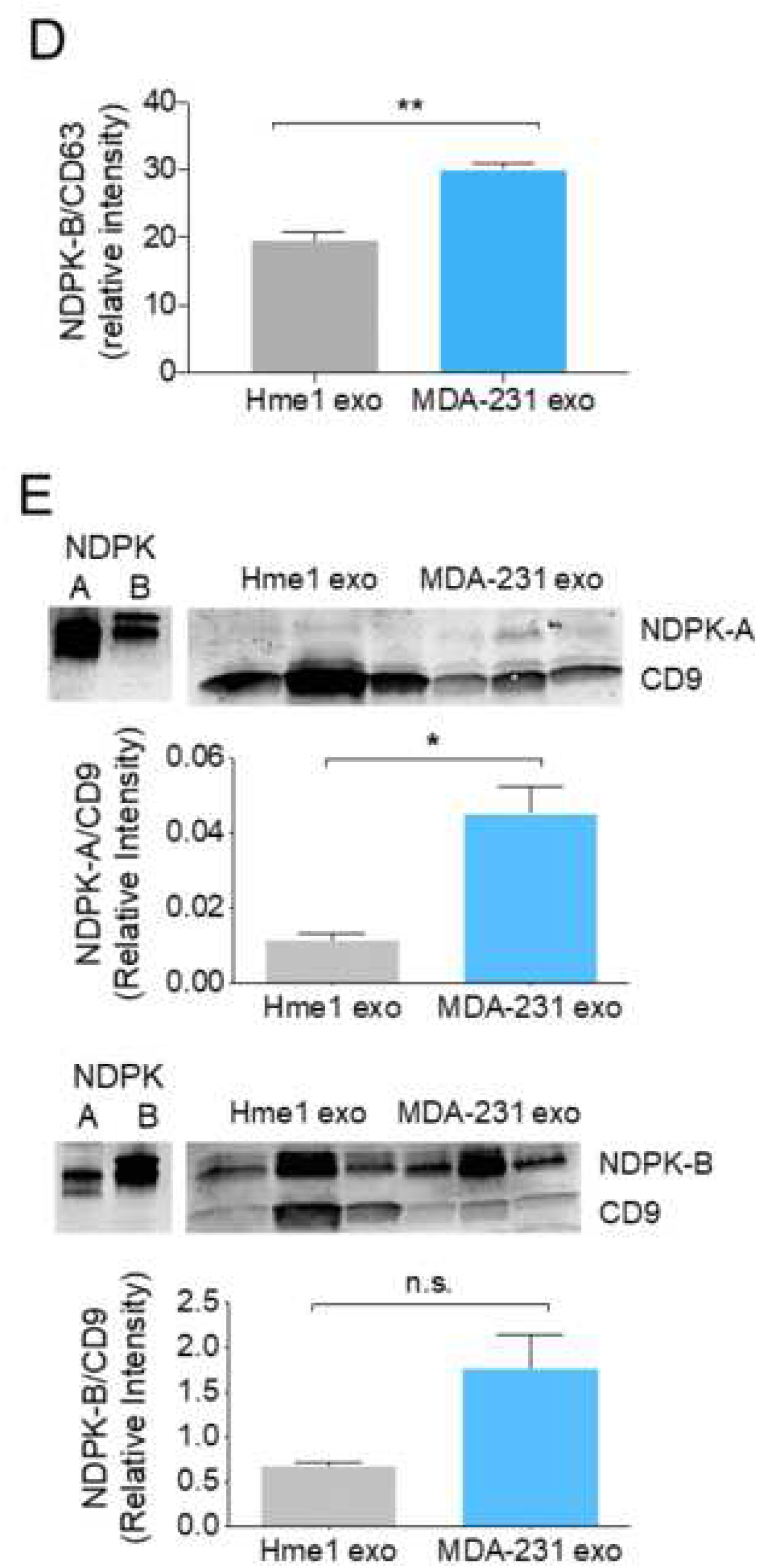
Non-tumorigenic breast epithelial cells and MDA-MB-231 breast cancer cells release EVs carrying NDPK-A and NDPK-B. *(A)* Flow cytometry analysis of HME1 and MDA-MB-231 EVs adsorbed onto latex beads and immuno-labeled for CD63 and *(B)* NDPK-A and NDPK-B. *(C)* Flow cytometry analysis of non-adsorbed HME1 and MDA-MB-231 EVs labeled for surface expression of CD63 and NDPK-A/B. Plots show side scatter area (y-axis) versus PE-fluorescence area (x-axis) on a log scale. *(D)* Quantitation of positive NDPK-B labeling in HME1 and MDA-MB-231 EVs normalized to CD63 intensity and reflecting background subtraction of respective isotype control. n = 3. *(E)* Western blot analysis of NDPK-A and NDPK-B expression normalized to CD9 in HME1 and MDA-MB-231 EV lysates. Cross-reactivity of antibodies was confirmed using recombinant NDPK-A/B protein. Mean ± S.E.M; two-tailed Student’s t-test, n = 3.

### MDA-MB-231 EVs are enriched in NDPK phosphotransferase activity

To determine whether elevated NDPK-B expression in EVs was correlated with higher NDPK phosphotransferase function, we performed a luciferin-luciferase ATP activity assay to measure NDPK-B phosphotransferase activity. Phosphotransferase activity was significantly higher in lysates compared to whole EVs, supporting the previous observation that NDPK is primarily associated with the intraluminal compartment. Additionally, MDA-MB-231 EV lysates demonstrated approximately three-fold higher transphosphorylase activity compared to HME1 lysates when assayed at equivalent concentrations (Figure 3A). We next tested whether ellagic acid (EA), a known inhibitor of NDPK transphosphorylase activity, could decrease the previously observed activity. Addition of EA reduced transphosphorylase activity of EVs in a dose-dependent manner, with a 10μM concentration reducing activity by two- to three-fold (Figure 3B). The P2Y1 receptor antagonist MRS2179 was not examined in this experiment as the products of NDPK activity are expected to act on the P2 receptor, not on the enzyme per se. We further confirmed these properties in ExoQuick-TC purified EVs (Supplemental Fig. 2).

**Figure 3.**
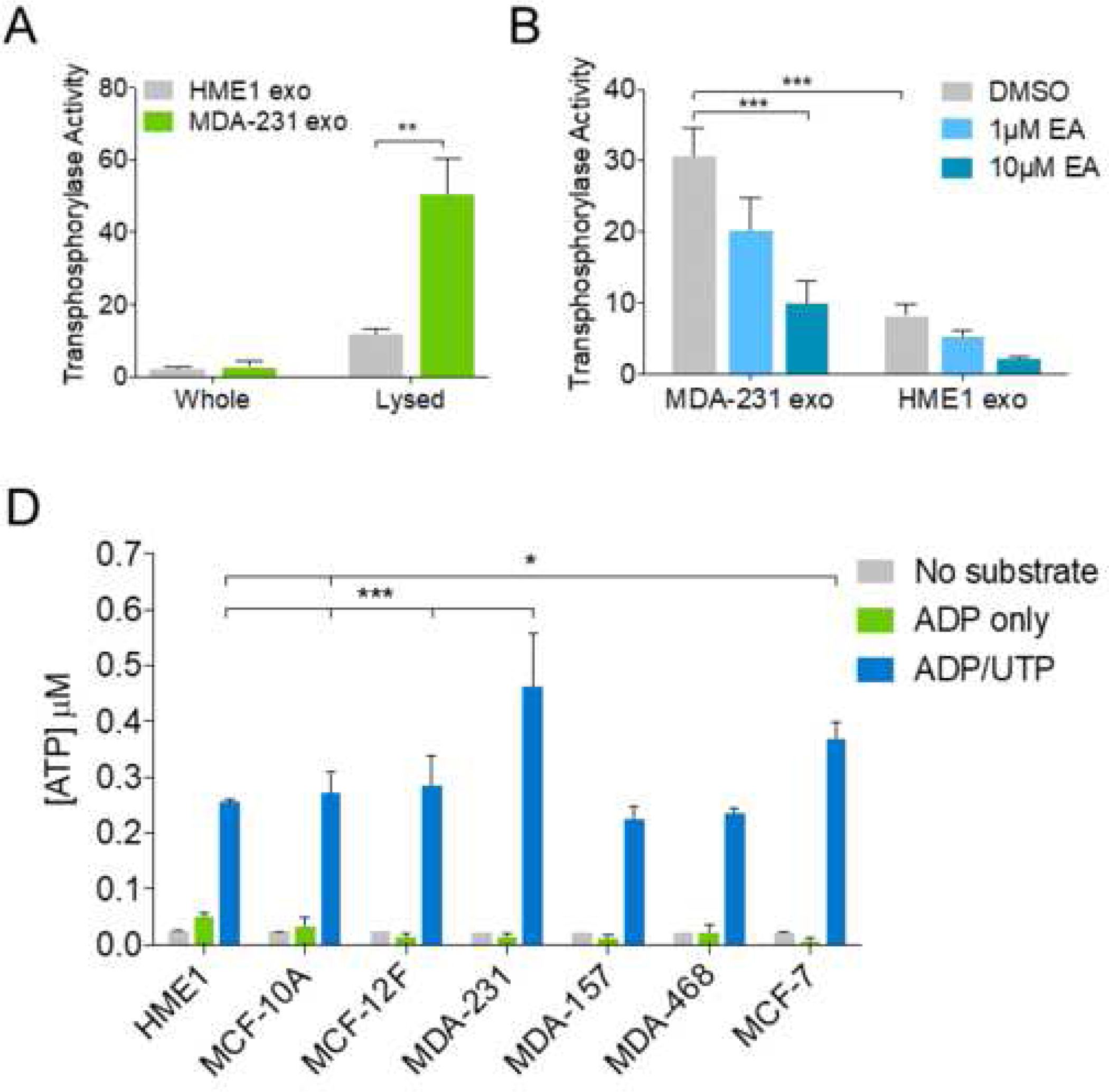
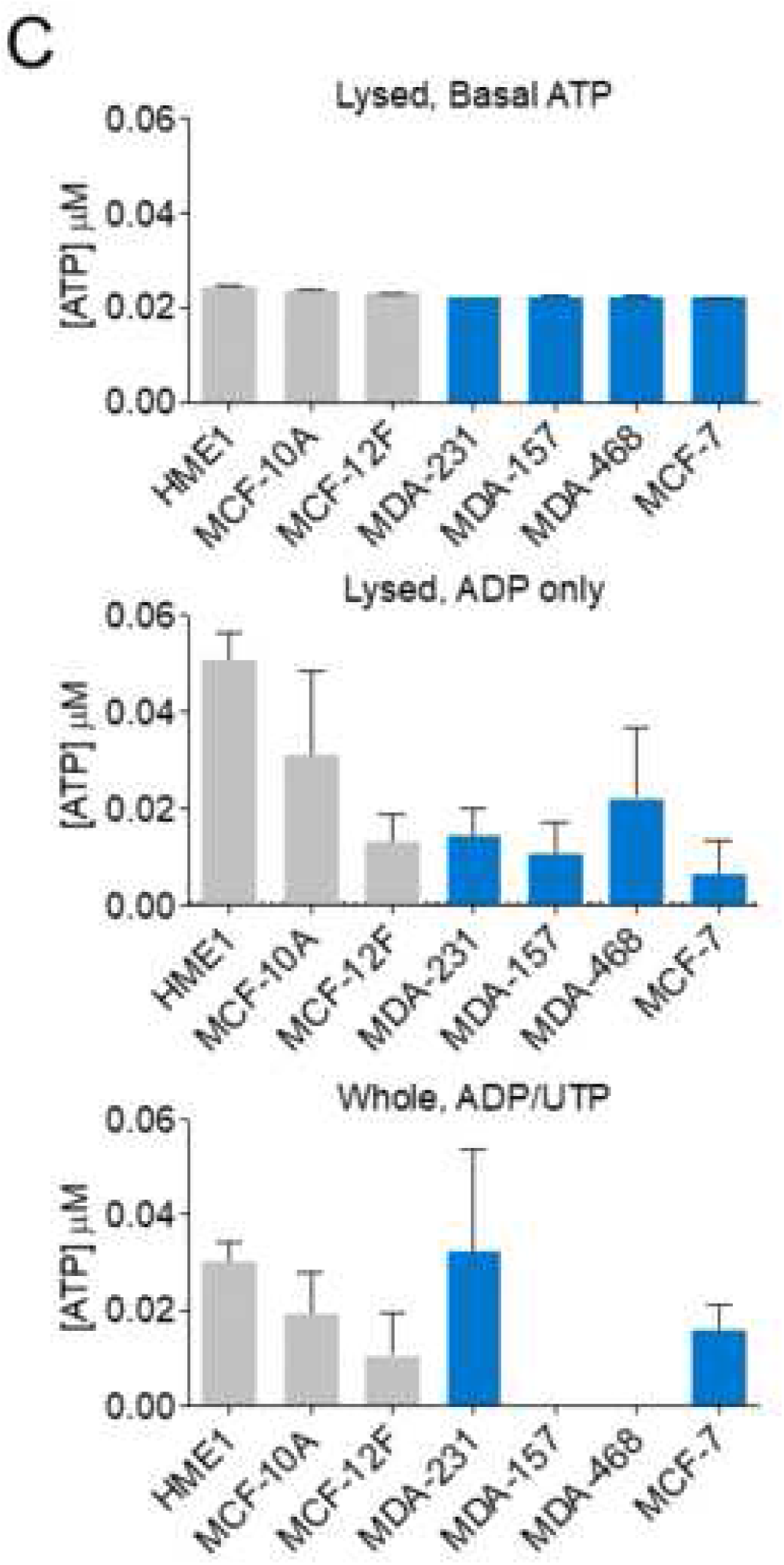
MDA-MB-231 EVs are enriched in NDPK transphosphorylase activity. *(A)* Measurement of NDPK transphosphorylase activity in whole and lysed EVs secreted by HME1 and MDA-MB-231 cells. n = 3 biological replicates. Mean ± S.E.M. \*\**p* < 0.05 by two-tailed Student’s t-test. *(B)* Measurement of NDPK transphosphorylase activity in MDA-MB-231 and HME1 EVs treated with ellagic acid (EA) or with DMSO vehicle. n = 3 biological replicates. Mean ± S.E.M. \*\*\**p* < 0.001 by two-way ANOVA. *(C)* Basal ATP production and ADP phosphorylating activity of EV lysates was measured, in addition to NDPK transphosphorylase activity of whole EVs. *(D)* EVs derived from multiple non-transformed and breast cancer cell lines were incubated with ADP/UTP substrate, ADP only, or no substrate and ATP production was measured. Mean ± S.E.M. \**p* < 0.05, \*\*\**p* < 0.001 by two-way ANOVA.

To test the robustness of these results, we evaluated NDPK phosphotransferase activity in EVs from two additional non-transformed human breast epithelial cell lines (MCF-10A and MCF-12F) and from three additional breast cancer cell lines (MDA-MB-157, MDA-MB-468, and MCF-7). NDPK phosphotransferase activity was highest in MDA-MB-231 and MCF-7 EVs, however no appreciable difference was observed between MDA-MB-468 and MDA-MB-157 EVs compared to those of non-transformed cells (Figure 3C). Moreover, ATP generation was predominantly attributed to NDPK activity, as both basal ATP production and adenylate kinase activity were ten times lower than NDPK activity in all EV populations (Figure 3D).

### NDPK inhibition and P2Y1 receptor antagonism reverse the pro-migratory effect of MDA-MB-231 EVs on vascular endothelial cells

To investigate the functional role of MDA-MB-231 EVs on vascular endothelium, we assayed migration as an indicator of endothelial cell activation, remodeling, and angiogenesis. We evaluated the effect of three concentrations of EVs on the migration of human umbilical vein endothelial cells (HUVECs). At two of the three concentrations tested, EVs derived from MDA-MB-231 cells demonstrated greater migratory-initiating capability compared to those of HME1 cells (Figure 4A). Treatment of HUVECs with MDA-MB-231 EVs enhanced migration after 24 hours compared to non-treated cells in low serum conditions. Inhibition of EV-associated NDPK transphosphorylase activity with EA significantly attenuated cell migration. As NDPK is known to activate downstream P2Y1 receptor signaling through the generation of extracellular ADP/ATP, we measured the effect of EV-mediated cell migration following blockade of the P2Y1 receptor. Delivery of the P2Y1 receptor antagonists MRS2179, MRS2279, and MRS2500 abrogated EV-mediated endothelial cell migration (Figure 4B). The robustness of these events were further tested in primary VCAM-1^+^ murine lung endothelial cells (MLECs). Consistent with the previous observations, MDA-MB-231 EVs stimulated migration of MLECs and combination treatment with EA and MRS2179 reversed these events (Figure 4C).

**Figure 4.**
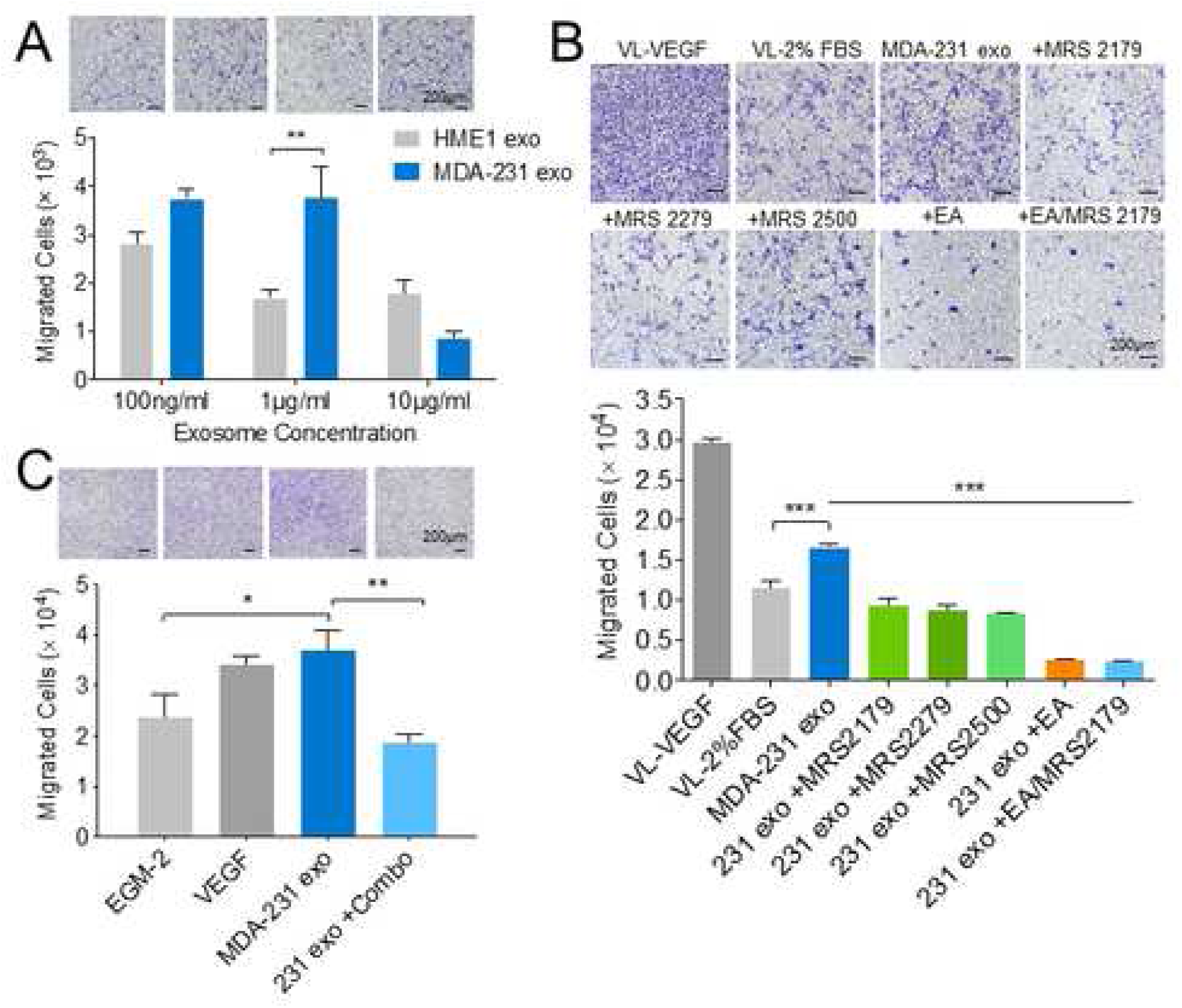
NDPK inhibition and P2Y1 receptor antagonism reverse MDA-MB-231 EV-mediated vascular endothelial cell migration. *(A)* Quantitation of HUVEC migration through collagen-coated transwells following 24-hour treatment with MDA-MB-231 EVs or HME1 EVs. Scale 200 μm. n = 3. Mean ± S.E.M. \*\**p* < 0.01 by nonparametric two-tailed Student’s t-test. *(B)* HUVEC migration through collagen-coated transwells following 24-hour treatment with complete growth medium (VL-VEGF), low serum (VL-2% FBS), MDA-MB-231 EVs (MDA-231 exo) with and without MRS2500, MRS2270, MRS2179, ellagic acid (EA), and ellagic acid with MRS2179. Scale 200 μm. n = 3. Mean ± S.E.M. \*\*\**P* < 0.001 by one-way ANOVA and Tukey post-test. *(C)* MLEC migration through collagen-coated transwells following 24-hour treatment with complete growth medium (EGM-2), VEGF, MDA-MB-231 EVs, and EVs with ellagic acid and MRS2179 (Combo). Scale 200 μm. n = 3. Mean ± S.E.M. \**p* < 0.05, \*\**p* < 0.01 by one-way ANOVA and Tukey post-test.

### NDPK inhibition rescues MDA-MB-231 EV-mediated permeabilization of vascular endothelial cell monolayers

As purinergic signaling has been implicated in vascular permeability leading to CTC dissemination, we next assayed the effect of varying MDA-MB-231 EV concentrations on endothelial cell monolayer integrity using a modified Boyden chamber approach (Supplemental Fig. 3). Treatment of human lung microvascular endothelial cells (HLMVECs) with MDA-MB-231 EVs increased monolayer permeabilization to FITC-labeled dextran compared to treatment with HME1 EVs and complete growth medium alone. Treatment with EA significantly reduced MDA-MB-231 EV-mediated permeabilization, whereas treatment with MRS2179 elicited a more moderate effect (Figure 5A). Similar results were observed in HUVECs following treatment with MDA-MB-231 EVs and EA. However, administration of MRS2179 alone or in combination with EA demonstrated no appreciable effect on reversing MDA-MB-231 EV-mediated HUVEC permeabilization (Figure 5B). These observations were consistent with a marked loss in monolayer integrity and junction-associated proteins ZO-1 and β-catenin. In addition, EA was shown to localize to the cell membrane, suggesting an intracellular membrane-associated mechanism for junction regulation (Figure 5C).

**Figure 5.**
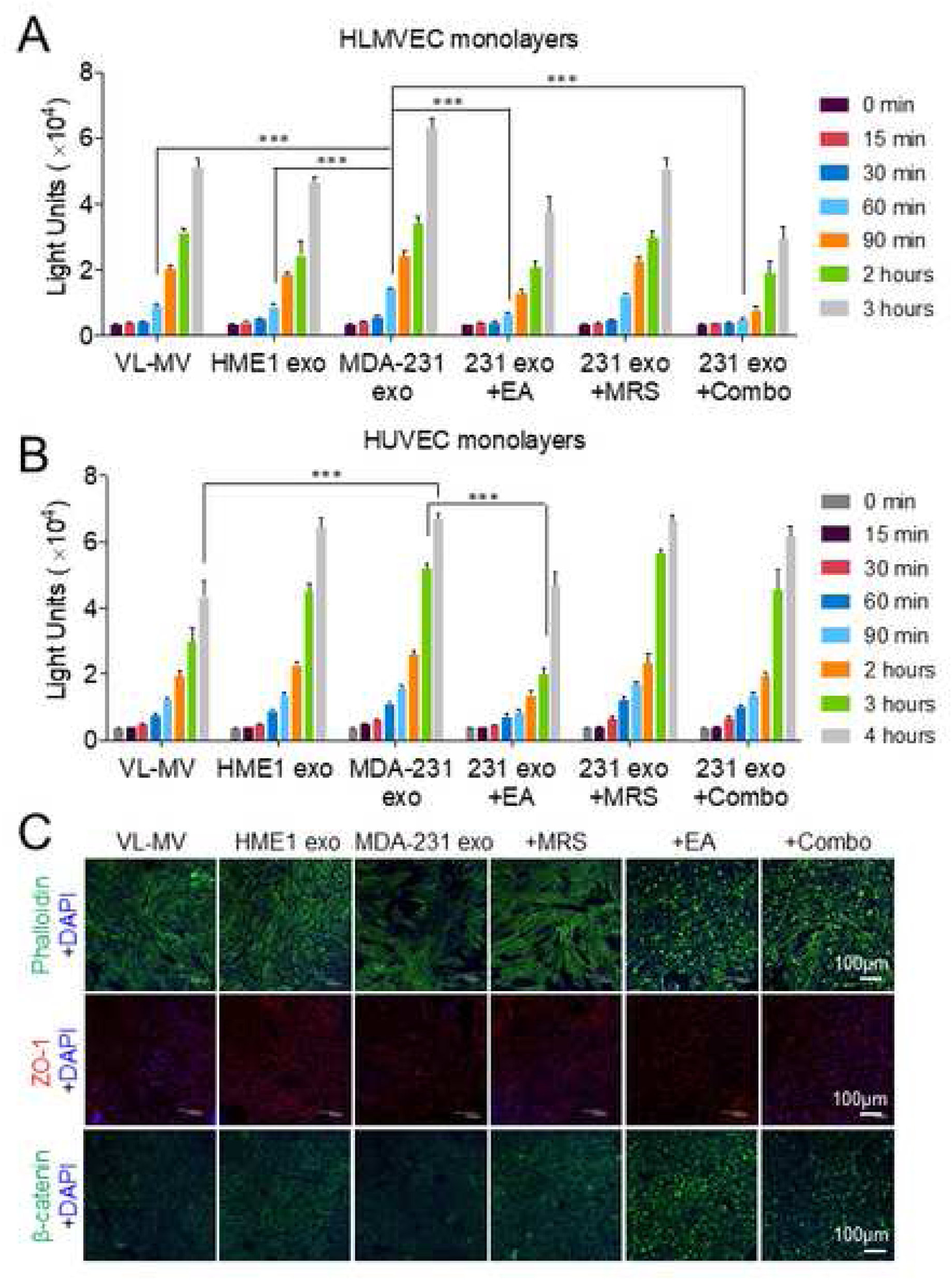
NDPK inhibition ameliorates endothelial monolayer permeabilization by MDA-MB-231 EVs. *(A)* Permeabilization of HLMVEC and *(B)* HUVEC monolayers to FITC-dextran was measured using Matrigel-coated tranwell chambers. Cells were treated with complete growth medium (VL-MV or VL-VEGF) with and without HME1 EVs, MDA-MB-231 EVs, and MDA-MB-231 EVs with MRS2179, EA, or a combination of both drugs. Intensity of FITC-dextran in top chambers was measured at indicated time points as an indicator of enhanced permeability. n = 6. Mean ± S.E.M. \*\*\**P* < 0.001 by one-way ANOVA and Tukey post-test. *(C)* CLSM images of HLMVEC monolayers immuno-stained for actin, ZO-1 and β-catenin expression. All CLSM laser acquisitions were kept consistent, except for bottom two images on the right, where 488 nm laser was enhanced to better visualize cells in these two treatment conditions. Cells were counterstained with DAPI. Scale 100 μm.

### MDA-MB-231 EVs enhance vascular leakage in the lung and NDPK inhibition or P2Y1 receptor antagonism ameliorates this effect

We further evaluated the ability of MDA-MB-231 EVs to disrupt *in vivo* vascular integrity. Treatment duration was determined by following previous studies that identified PMN formation after three weeks of EV treatment ^[8,9]^. EV exposure was subsequently prolonged to eight weeks to better mimic physiological metastasis. Following three- or eight-week treatment with MDA-MB-231 EVs, mice were injected systemically with Evans Blue dye (EBD) and lungs were imaged to visualize extravasated dye. Three-week MDA-MB-231 EV treated mice demonstrated enhanced pulmonary vascular leakage (Figure 6A). Similarly, the perfused lungs of eight-week MDA-MB-231 EV treated mice demonstrated higher levels of EBD signal compared to PBS treatment. Treatment with EA or MRS2179 individually reduced EBD signal in lung tissues, however combination treatment showed no appreciable effect on EBD infiltration (Figure 6B). We further measured the absorbance of extracted EBD from perfused lungs. Consistent with the previous observations, EA or MRS2179 treatment reversed MDA-MB-231 EV-mediated vascular permeabilization, as indicated by a decreased amount of extravasated dye in lung tissue (Figure 6C).

**Figure 6.**
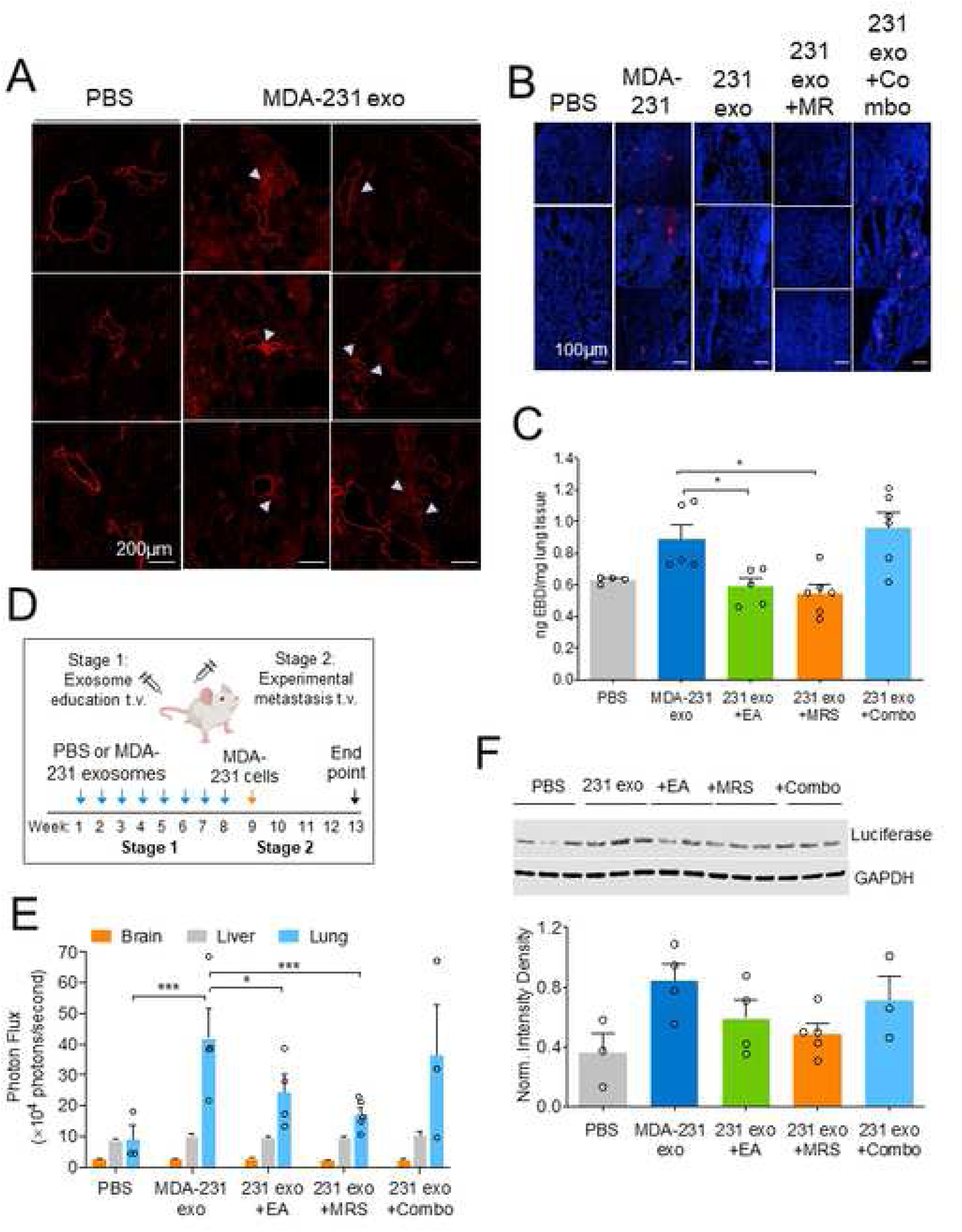
Inhibition of NDPK phosphotransferase activity or P2Y1 receptor blockade blunts MDA-MB-231 EV-mediated vascular leakage and experimental lung metastasis. *(A)* CLSM images of lungs from mice treated for three weeks with MDA-MB-231 EVs or PBS vehicle and systemically injected with EBD emitting in the far red spectrum. Arrows indicate regions of vascular leakage. n = 3 per group. Scale 200 μm. *(B)* Representative CLSM images of lungs from mice treated for 8 weeks with PBS or MDA-MB-231 EVs in combination with drug inhibitors. Mice were systemically injected with EBD (red) and perfused to clear EBD from lung vasculature. Lung sections were counterstained with DAPI. Scale 100 μm. *(C)* EBD was biochemically extracted from 8-week treated perfused mouse lungs and quantified by measuring absorbance at 610 nm. n = 4-6 per group. Mean ± S.E.M. \**P* < 0.05 by one-way ANOVA and Tukey post-test. *(D)* Visual summary of experimental metastasis study design. *(E)* The brain, liver, and lungs of 8-week treated mice were imaged *ex vivo* for bioluminescence intensity to detect the presence of experimental MDA-MB-231-Luc^+^ metastases. n = 3-5 per group. Mean ± S.E.M. \**p* < 0.05, \*\*\**p* < 0.001 by two-way ANOVA. *(F)* Western blot analysis of luciferase expression relative to GAPDH in the right lung lobe of 8-week treated mice. n = 3-5 per group. Data n.s. by one-way ANOVA.

NDPK-mediated activation of P2Y1/2 receptors on endothelium is known to stimulate pro-inflammatory and vaso-dilative factors, including nitric oxide synthase (NOS) and cyclooxygenase (COX). We mimicked the conditions used in previous *in vitro* assays and measured NOS and COX expression following delivery of EVs with a high dosage of inhibitors for three hours, or with a lower drug dosage for 24 hours. Comparison between these conditions provides a broader understanding of the immediate and long-term signaling effects mediated by EVs. Three and 24-hour treatment of HUVECs with MDA-MB-231 EVs led to a moderate, non-significant increase in eNOS expression, whereas treatment with a high dosage of MRS2179 (100 μM) attenuated this effect. Additionally, 24-hour treatment with a lower dosage of EA (10 μM) or combination drug treatment significantly decreased COX-1, COX-2, and iNOS expression, whereas three-hour treatment with a higher dosage of both drugs (100 μM) induced a three-fold increase in COX-2 expression (Supplemental Fig. 4).

### Attenuation of eNDPK activity or P2Y1 receptor activation blunts the development of lung metastases in an experimental metastasis model

We next considered whether enhanced vascular permeability promotes metastasis of CTCs to the lung. Following eight-week treatment, mice were injected systemically with MDA-MB-231-Luc+ cells expressing CD63-GFP and metastasis to the lungs, liver, and brain was evaluated 30 days later (Figure 6D). While bioluminescence was detected in all mouse lungs, no signal was detected in the liver or brain, indicating the absence of overt metastases in these tissues (Supplemental Fig. 5). Lungs from mice treated with MDA-MB-231 EVs displayed higher bioluminescence intensity and luciferase protein expression compared to those of the control group, suggesting a greater presence of colonized MDA-MB-231 cells. Treatment with EA or MRS2179 reduced lung bioluminescence intensity and luciferase expression, however collective administration of these drugs appeared to elicit no beneficial effect (Figure 6E and F). These observations were concomitant with a slight increase in NDPK-A/B levels in three-week treated experimental mice, however no difference was observed following prolonged MDA-MB-231 EV treatment (Supplemental Fig. 6).

Intriguingly, prolonged MDA-MB-231 EV exposure leading to vascular remodeling and metastatic outgrowth was not associated with a change in circulating angiogenic cytokines that have been previously suggested in PMN formation, including CXCL12, EGF, FGF-2, PLGF, and VEGF-A. However, MDA-MB-231 EV induced significant fluctuations in host production of immuno-modulatory cytokines, including suppression of TNF-α, IL-6, IL-10, and induction of the chemokines CCL2, CCL3, and CCL5 (Supplemental Fig. 7).

### Proteomic analysis of the lung following MDA-MB-231 EV treatment identifies purinergic events known to support pre-metastatic niche formation

To identify potential changes in the lung proteome that led to the previous functional observations, we performed mass spectrometry on three-week treated lungs bearing tandem mass tags. We identified 36 proteins that were differentially expressed between control and experimental cohorts at the *p* < 0.05 significance level with log2 fold-change > 0.5 (Figure 7A). A Principal Component Analysis of statistically significant proteins with raw p-values of *p* < 0.01 shows a distinct separation between control and experimental groups, with the first component capturing 65% of the experimental variance (Figure 7B).

**Figure 7.**
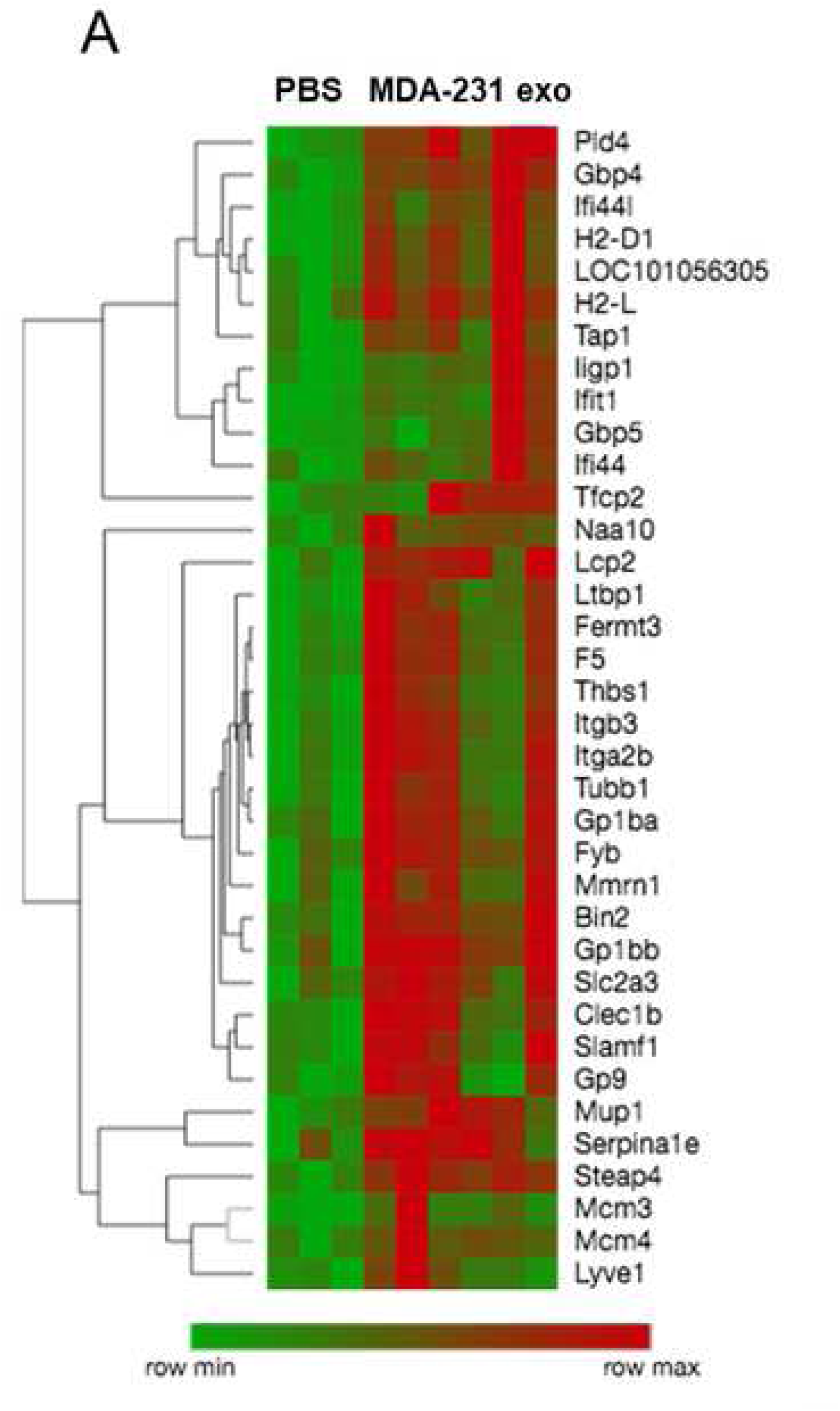
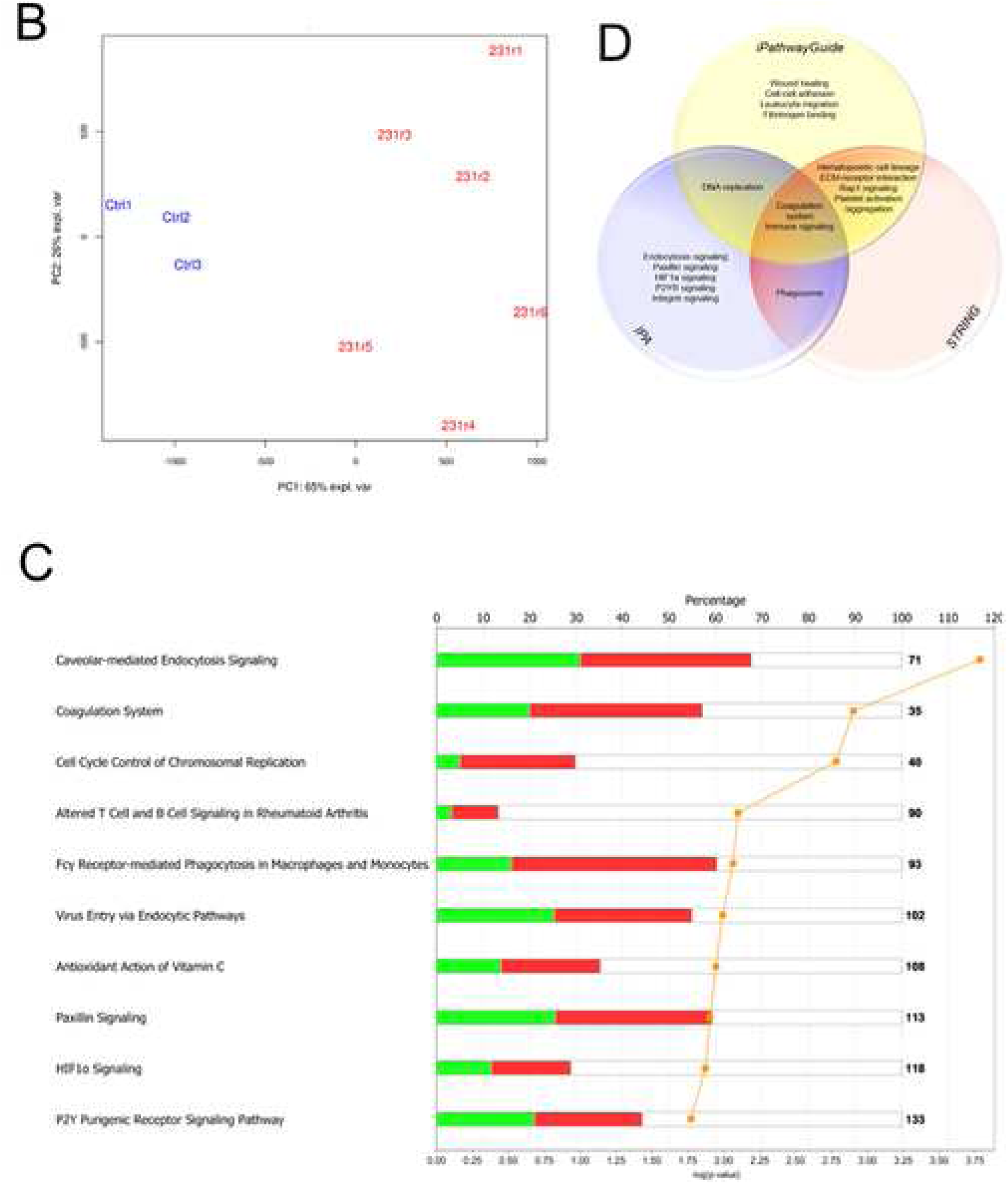
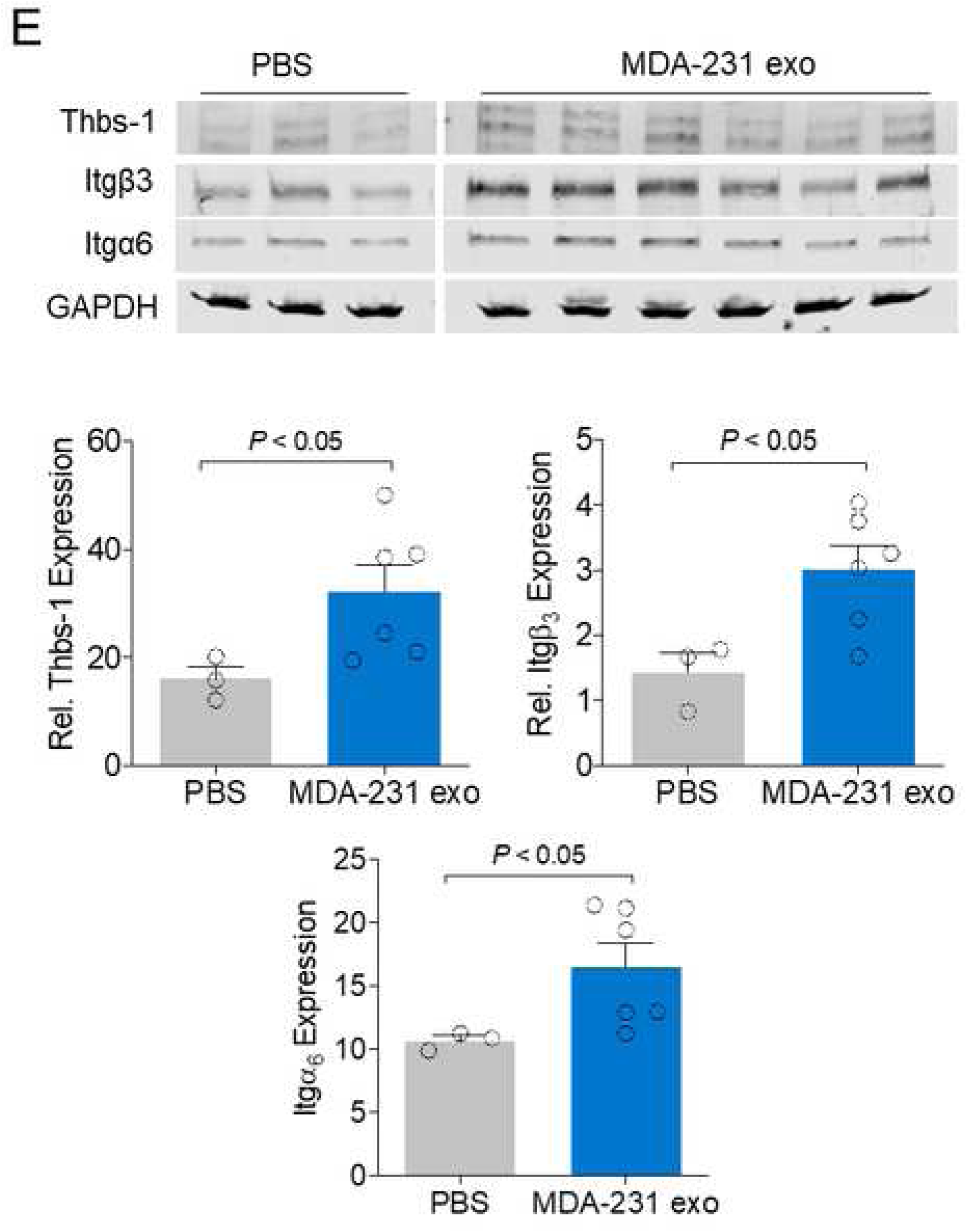
MDA-MB-231 EVs induce proteomic transformation in the lung and activate purinergic events associated with pre-metastatic niche formation. Mass spectrometry was performed on TMT-labeled lungs from mice treated for three weeks with PBS (n = 3) or MDA-MB-231 EVs (n = 6). *(A)* Heatmap with hierarchical clustering of differentially expressed proteins at the *p* < 0.05 significance level with log2 fold-change above 0.5. *(B)* Principal component analysis of top 107 differentially expressed proteins with raw *p-*values < 0.01. n = 3 for PBS treated mice (Ctrl) and n = 6 for MDA-MB-231 EV treated mice (231r). *(C)* Top ten canonical pathways perturbed by MDA-MB-231 EV treatment, as identified by IPA analysis. *(D)* Venn diagram of significant pathways, biological processes, and molecular functions identified by IPA, iPG, and STRING software. FDR correction was applied at the protein dataset level and during pathway analyses with significance defined at *p* < 0.05.

Pathway ontology analysis on differentially expressed proteins (*p* < 0.05) with log2 fold-change over 0.5 was performed using Ingenuity Pathway Analysis (IPA), iPathwayGuide (IPG), and STRING software. Proteins with the greatest log2 fold-change were identified and ranked (Supplemental Tables 1 and 2). Relevant canonical pathways identified by IPA include caveolar-mediated endocytosis, coagulation, hypoxia inducible factor (HIF-1α) signaling, and P2Y purinergic receptor signaling (Figure 7C). All three software tools identified perturbations to the coagulation system and immune response, whereas iPG and STRING identified the involvement of hematopoietic cell lineage, ECM-receptor interaction, and platelet activation and aggregation pathways. Pathways and processes unique to iPG include wound healing, cell adhesion, and fibrinogen binding (Figure 7D). Additionally, activated cellular components include the integral, internal, and external components of the plasma membrane, indicating internalization and/or membrane receptor interaction with EVs. The validity of this experiment was confirmed by western blot of select differentially expressed proteins identified by mass spectrometry (Figure 7E). Collectively, these analyses identify purinergic events known to support PMN formation.

To add further depth to these experiments, we assessed proteomic cargo associated with MDA-MB-231 EVs and EVs from the non-tumorigenic MCF-12F cell line. Significant differentially expressed proteins with over three-fold enrichment were identified in MCF-12F and MDA-MB-231 vesicles (Supplemental Fig. 8, Supplementary Tables 3 and 4). Processes associated with MDA-MB-231 EVs include cell migration, ECM disassembly, and positive regulation of angiogenesis. In support of the previous observations, these findings suggest that EV cargoes reflect parent cell lineage and propagate their malignant features.

## Discussion

Since its identification 30 years ago as the first novel metastasis suppressor gene ^[10]^, Nm23 and its prognostic value in cancer remain controversial. Reduced Nm23-H1 mRNA and NDPK expression levels have long been reported in solid tumors, and loss of Nm23 correlates with disease progression and metastasis in several cancers. Investigations by our lab and others into extracellular Nm23 illuminate a diverging plotline for its role in oncogenesis. Among these, elevated Nm23 expression was reported in sera of patients with breast cancer ^[31]^, colorectal cancer ^[38,39]^, neuroblastoma ^[40]^, renal carcinoma ^[41]^, and hematological malignancies ^[42]^. Metastatic breast cancer cells of human and murine origin secrete elevated levels of NDPK-A and -B isoforms compared to non-tumorigenic breast cells, suggesting a functional role for secreted NDPK in metastasis ^[43,44]^. However, the mechanism by which NDPK-A/B is elaborated into the extracellular milieu remains unclear. Here, we showed that the highly metastatic and triple negative MDA-MB-231 cell line propagates functional NDPK-B phosphotransferase activity through the secretion of EVs.

We corroborate findings by several other groups showing pro-angiogenic and endothelium-remodeling properties associated with tumor-secreted EVs ^[45,47]^. In accordance with the Nucleotide Axis Hypothesis first proposed by Buxton and colleagues ^[21]^, we present a role for exosomal NDPK in mediating P2Y1 receptor-dependent endothelial cell activation and remodeling. Interestingly, our observations suggest that dual inhibition of NDPK and P2Y1 receptor activation diminishes the beneficial effects observed when drug compounds are delivered individually. Further work is required to elucidate whether these events are attributed to the observed induction of COX-2 following high dosage treatment in endothelial cells, or whether P2Y2R and other purinoreceptors exert a compensatory effect. Indeed, activation of P2Y2 receptor by eATP has been shown to promote PMN formation ^[33]^ and enhance the invasiveness of breast cancer cells via crosstalk with endothelium ^[24]^. Thus, blocking dual P2Y1/2 receptor activation may potentiate their individual anti-metastatic effects.

Cancer-derived exosomes have been shown to be enriched in a variety of both pro-angiogenic and angio-static factors. While we did not presently explore the contributions of the latter, it is well established in the literature that such antagonistic signaling factors may influence endothelial cell activation and suppression. Based on our observations, the pro-migratory effect induced by EV NDPK may be opposed by other anti-angiogenic factors at higher EV concentrations. For instance, our identification of thrombospondin-1 as a significantly enriched factor in breast cancer EVs lends evidence to support such antagonistic signaling. Future application of more specific NDPK silencing approaches, such as the delivery of shRNA and siRNA, would enable a further dissection of these signaling mechanisms.

As NDPK phosphotransferase activity was predominantly associated with the intraluminal exosome compartment, there remains the question of how NDPK mediates extracellular nucleotide homeostasis. One possibility exists whereby exosomal NDPK is released into the cytosol of the recipient cell following direct membrane fusion. Previous reports identify a function for membrane-associated NDPK as a nucleotide reservoir and as a required component for caveolin and G protein-mediated VEGFR-2 activation ^[47,48]^. Direct exosome fusion may also facilitate ectopic NDPK expression on the recipient cell surface (Figure 8). Such a mechanism is supported by previous reports of NDPK localizing to caveolae and binding to the membrane ^[49,50]^. As these membranous components are known to be recycled during exosome biogenesis, it thus remains a possibility that NDPK associates with the exosome membrane and is transferred to recipient cell membranes. Once ectopically expressed, NDPK can propagate its phosphotransferase activity to activate adjacent P2Y1/2 receptors co-expressed on endothelium. This proposed mechanism is congruent with previous observations by our group that NDPK is ectopically expressed on vascular endothelium.

**Figure 8.**
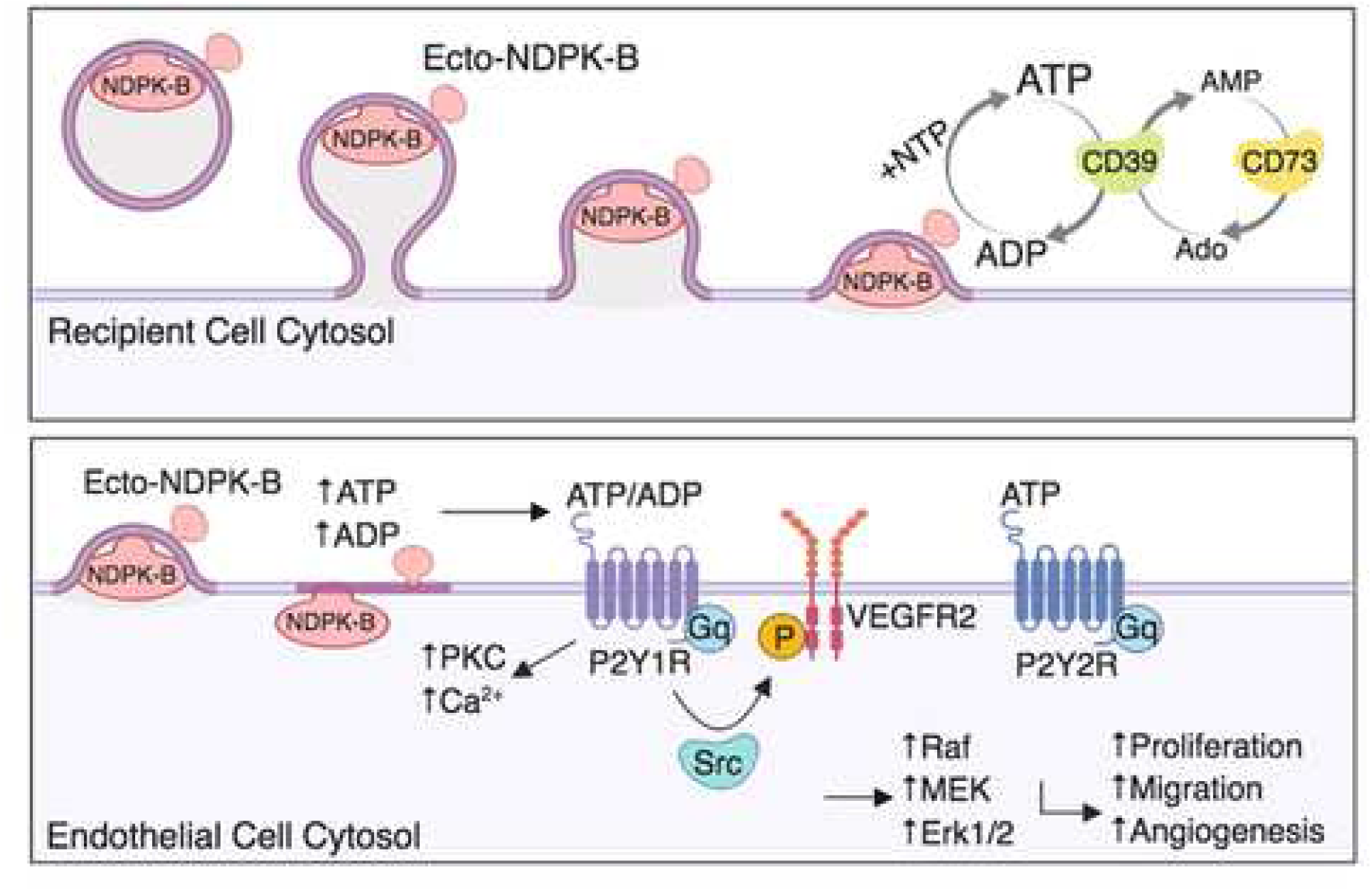
Proposed mechanism of extracellular vesicle-mediated NDPK signaling in endothelial cell activation. Ectopic NDPK-B maintains extracellular ATP and ADP pools by catalyzing phospho-transfer between NTP and NDP substrates. Extracellular ATP and ADP activate the P2Y1 purinergic receptor expressed on vascular endothelium to stimulate known signaling pathways involved endothelial cell activation, remodeling, and angiogenesis.

We identified MDA-MB-231 EV-mediated perturbations to the P2Y and HIF-1α signaling pathways, lending evidence to support purinergic regulation of HIF-1α–LOX signaling in PMN formation ^[33]^. Of further interest were pathways involved in platelet activation, aggregation, and wound healing, as these responses are mediated by purinergic signaling and are implicated in PMN formation and metastasis ^[51,54]^. These analyses suggest that MDA-MB-231 EVs activate platelets, potentially through eNDPK-mediated activation of P2Y1 and P2Y12 receptors expressed on platelets. Further investigation into the role of eNDPK in modulating these purinergic pathways may reveal additional downstream effectors that can be targeted therapeutically to blunt metastatic progression.

Finally, our observations raise a previously unvisited question concerning the homeostatic dynamic of EV-mediated NDPK signaling. In addition to NDPK, several other ectoenzymes mediate purine metabolism and homeostasis in the tumor microenvironment, including CD39, CD73, adenosine deaminase and adenylate kinase. The importance of their collective roles in the tumor microenvironment is undermined by the fact that both adenosine and ATP are present at remarkably high levels in tumor interstitium compared to in non-tumor bearing tissues ^[55, 56]^. Moreover, adenosine and ATP mediated activation of purinergic receptors has been reported to play a role in modulating immune cell function, angiogenesis, and the inflammatory response ^[57]^. While the studies presented here sought to investigate EV-mediated NDPK signaling in isolation, further work examining the relationship between NDPK and other ectonucleotidases will shed additional light on homeostatic signaling within the tumor microenvironment.

## Supporting information

All 8 supplemental figures and 4 supplemental tables

## Acknowledgments

We would like to acknowledge the following individuals for their assistance with proteomics data acquisition and analysis: Dr. Juli Petereit, Dr. Craig Ulrich, Dr. Karen Schlauch, Dr. David Quilici, and Rebekah Woolsey.

## Authors’ Contributions

SD, SN, and ILOB designed experiments and contributed to data analysis and interpretation. SD, SN, and AEB conducted experiments. SD and ILOB contributed to writing the manuscript. All authors have read and approved the final manuscript.

## COMPLIANCE WITH ETHICAL STANDARDS

### Use of Animals in Research

The Institutional Animal Care and Use Committee of the University of Nevada, Reno in compliance with the US Department of Agriculture Animal Welfare Regulations, International AAALAC standards, and the Public Health Service Policy on Humane Care and Use of Laboratory Animals approved use of mice in our experiments.

### Informed Consent

Human subjects were not participants in this work.

## Supplemental Table and Figure Legends

**Supplemental Table 1. Top 20 up-regulated lung proteins after MDA-MB-231 EV treatment.** Mice were treated for three-weeks with MDA-MB-231 EVs or PBS vehicle and TMT-labeled lungs were submitted for mass spectrometry analysis. Significantly enriched proteins in MDA-MB-231 EV-treated lungs with raw p-values less than 0.05 were ranked by log2 fold-change.

**Supplemental Table 2. Top 20 down-regulated lung proteins after MDA-MB-231 EV treatment.** Mice were treated for three-weeks with MDA-MB-231 EVs or PBS vehicle and TMT-labeled lungs were submitted for mass spectrometry analysis. Significantly down-regulated proteins in MDA-MB-231 EV-treated lungs with raw p-values less than 0.05 were ranked by log2 fold-change.

**Supplemental Table 3. Top 20 proteins with greatest fold enrichment in MDA-MB-231 EVs.** Significantly enriched proteins in MDA-MB-231 EVs with adjusted p-values less than 0.01 were ranked by log2 fold-change.

**Supplemental Table 4. Top 20 proteins with greatest fold enrichment in MCF-12F EVs.** Significantly enriched proteins in MCF-12F EVs with adjusted p-values less than 0.01 were ranked by log2 fold-change.

**Supplemental Figure 1. Characterization of ExoQuick-TC-isolated HME1 and MDA-MB-231 EVs by TEM, CLSM, and flow cytometry**. *(A)* TEM images of EVs isolated from MDA-MB-231 cells. Scale 20 and 100 nm. *(B)* MDA-MB-231 and HME1 EVs immuno-gold labeled for tetraspanins, respectively. Scale 100 nm. *(C)* CLSM images of MDA-MB-231 cells and purified MDA-MB-231 EVs expressing GFP or RFP-labeled CD63. Scale 100 μm. *(D)* Flow cytometry analysis of CD63-labeled MDA-MB-231 and HME1 EV populations.

**Supplemental Figure 2. MDA-MB-231 EVs isolated by ExoQuick-TC are enriched in NDPK-A/B expression and transphosphorylase activity.** *(A)* Super-resolution image of a single MDA-MB-231-CD63-GFP expressing cell (left) and a representative MDA-MB-231 EV (right) immuno-stained for NDPK-B (red) expression. Scale 500 nm. *(B)* Flow cytometry analysis of MDA-MB-231 and HME1 EV populations immuno-labeled for NDPK-A expression. *(C)* Western blot analysis of lysed MDA-MB-231 and HME1 EVs and relative quantitation of NDPK-A/B expression. Recombinant NDPK-A, NDPK-B, and MDA-MB-231 cell lysate (CL) are also shown for comparison. n = 3. Mean ± S.D. \**p* < 0.05 by two-tailed Student’s t-test. *(D)* Measurement of NDPK activity in MDA-MB-231 and HME1 EVs using an ATP transphosphorylase activity assay. n = 3. Mean ± S.D. \**p* < 0.05 by two-tailed Student’s t-test. *(E)* Measurement of transphosphorylase activity in MDA-MB-231 and HME1 EVs treated with ellagic acid, an inhibitor of NDPK, or with DMSO vehicle. n = 3. Mean ± S.D. \**p* < 0.05, \*\**p* < 0.01, \*\*\**p* < 0.001 by two-way ANOVA.

**Supplemental Figure 3. The effect of varying MDA-MB-231 EV concentrations on endothelial monolayer permeabilization to FITC-dextran.** HUVEC monolayers seeded in Matrigel-coated tranwell chambers were treated with MDA-MB-231 EVs in complete growth medium (VL-VEGF). Intensity of FITC-dextran permeabilization was measured at indicated time points as a measure of enhanced permeability. n = 3. Mean ± S.E.M. **p < 0.01 and \*\*\**p* < 0.001 by one-way ANOVA and Tukey post-test.

**Supplemental Figure 4. Expression of COX-1, COX-2, iNOS, and eNOS in three- and 24-hour treated HUVECs.** *(A)* Wes analysis of three-hour treated HUVECs with EVs (100 ng/ml) and high dose drug inhibitors (100 μM EA, MRS2179, or both). *(B)* Western blot analysis of 24-hour treated HUVECs with EVs (100 ng/ml) and a lower dose of drug inhibitors (10 μM EA, MRS2179, or both). n = 3. Mean ± S.E.M. \**p* < 0.05, \*\**p* < 0.01, and \*\*\**p* < 0.001 by one-way ANOVA and Tukey post-test.

**Supplemental Figure 5. Bioluminescence images of experimental metastases in brain, lung, and liver tissue from MDA-MB-231 EV treated mice.** Mice were treated for eight weeks with vehicle, MDA-MB-231 EVs, or MDA-MB-231 EVs with the P2Y1 receptor antagonist MRS2179, the NDPK inhibitor ellagic acid (EA), or a combination of both drugs (Combo). Following treatment, MDA-MB-231-Luc^+^ cells were injected by tail vein and development of metastases at select organs was evaluated by bioluminescence imaging 30 days later.

**Supplemental Figure 6. Western blot analysis of NDPK-A and NDPK-B expression in MDA-MB-231 EV treated mouse lungs.** *(A)* NDPK-A and NDPK-B expression in lung lysates of mice treated for three weeks with PBS (n = 3) or MDA-MB-231 EVS (n = 6). *(B)* NDPK-A expression in lung lysates of mice treated for eight weeks with the indicated treatments. Following eight weeks of treatment, mice were injected by tail vein with MDA-MB-231-Luc^+^ cells and lungs were evaluated 30 days following injection (n = 3).

**Supplemental Figure 7. Prolonged MDA-MB-231 EV exposure induces opposing actions on systemic cytokine production.** Circulating cytokines were evaluated in the sera of SCID mice that were treated for eight weeks with MDA-MB-231 EVs or PBS vehicle. n = 4 per group; * = *p* < 0.05, # = *p* = 0.05, ## = *p* = 0.06 by unpaired Student’s t-test. Values reflect means of log2 (fold change) relative to PBS control group.

**Supplemental Figure 8. Validation of mass spectrometry analysis by western blot of select differentially upregulated proteins in lungs of MDA-MB-231 EV-treated mice.** NCr/SCID mice were injected by tail vein three times a week for three weeks with PBS vehicle or MDA-MB-231 EVs (10 μg). Lung lysates were analyzed by western blot for the expression of differentially expressed proteins, as previously identified by mass spectrometry. n = 3, PBS treated. n = 6, MDA-MB-231 EV treated. Mean ± S.E.M. \**P* < 0.05 by two-tailed Mann-Whitney test.

**Supplemental Figure 9. Proteomic profiling of ExoQuick-TC-isolated MDA-MB-231 EVs reveals malignant features consistent with parent cell line.** *(A)* Heatmap with hierarchical clustering of significant differentially expressed (DE) proteins with over three-fold log2 expression difference between MDA-MB-231 EVs and MCF-12F EVs. n = 3 per group. *(B)* Significant DE proteins were mapped to biological processes associated with MDA-MB-231 EVs. Proteins with red nodes denote enrichment in MDA-MB-231 EVs, while green nodes represent proteins significantly upregulated in MCF-12F EVs.

## Supplemental Materials

### Expanded Methodology

#### Super-Resolution Microscopy

For Supplemental Figure 2, EVs were fixed onto fibronectin-coated glass slides in 4% PFA, then blocked in 5% BSA and incubated in 1:100 CPTC-NME2-3 primary antibody overnight (Developmental Studies Hybridoma Bank, Iowa City, IA). Slides were incubated in 1:100 AlexaFluor 680 secondary antibody (Thermo Fisher Scientific) and mounted onto depression slips. Slides were imaged on the Leica GSD super resolution microscope (Leica Microsystems Inc., Buffalo Grove, IL).

#### Transmission Electron Microscopy (TEM)

Briefly, EVs were diluted in 2% PFA and adsorbed onto Formvar-carbon coated nickel grids, washed with 50 mM glycine, and blocked with 0.5% bovine serum albumin (BSA). For immunolabeling shown in Supplemental Figure 1, grids were incubated with CD9, CD81, and CD63 antibodies (Biolegend) and secondary antibodies conjugated with 10 nm gold particles (Abcam, Cambridge, MA).

#### TMT-labeled Mass Spectrometry

Samples were processed according to Lundby et al. (2012) and mass tagged using a TMT 10-plex isobaric label kit (Thermo Fisher Scientific). Pooled TMT-labeled peptides were fractionated by basic pH reversed-phase fractionation on an Ultimate 3000 HPLC (Thermo Fisher Scientific). BPRP fractions were separated using an UltiMate 3000 RSLCnano system (Thermo Fisher Scientific). Mass spectral analysis was performed using an Orbitrap Fusion mass spectrometer (Thermo Fisher Scientific). TMT analysis was performed using an MS3 multi-notch approach ^[36]^. Data was analyzed using Sequest (Thermo Fisher Scientific, version v.27, rev. 11.) and Proteome Discoverer (Thermo Fisher Scientific, Version 2.1). A minimum coverage of two peptides was required for positive protein identification.

#### Protein Level Data Analysis

4,992 proteins with abundance values in all control samples and at least three experimental samples met our criteria for further study. The abundance data of these 4,992 proteins were normalized at the protein level, following the TMT Thermo-Fisher protocol and manual (Thermo Proteome Discoverer User Guide, Version 2.1), using the sum of the abundances of a pool of the nine samples included on the multiplex. Normalized data were log2-transformed to follow a normal distribution. Control and experimental abundance values were examined by histogram and quantile (Q-Q) plot to confirm a normal distribution. As each protein was examined as an independent entity and each protein’s cohorts included at least three values, the Shapiro-Wilk test was used to test for normality. 208 proteins did not pass the test for normality and were excluded from analyses. Student’s t-tests were performed on the normalized data of 4,784 proteins and a correction for the false discovery rate (FDR) was applied ^[37]^.

#### Pathway Analysis

iPathwayGuide (iPG) version 1711 (Advaita Corporation, Plymouth, MI), Ingenuity Pathway Analysis (IPA) (Qiagen, Redwood City, CA), and STRING database version 10.5 (STRING Consortium) were used to identify affected pathways. A cutoff criteria of *p* < 0.05 with log2 fold-change ≥ 0.5 for differentially expressed proteins was applied. Proteins with known symbols (UNIPROT) and their corresponding expression values were uploaded into each database and protein symbols were mapped to their corresponding genes. For IPA, canonical pathways most significant to the input data set were identified using the IPA library of canonical pathways. Significance was based on two parameters: (1) A ratio of the number of genes from the data set that map to the pathway divided by the total number of genes that map to the canonical pathway and (2) a *P* value calculated using Fisher’s exact test determining the probability that the association between the genes in the data set and the canonical pathway is due to chance alone. The log2 normalized expression values of the differentially expressed proteins were mapped using the open source online-tool Morpheus (Broad Institute).

#### Experimental Metastasis Study

For cytokine analysis presented in Supplemental Figure 7, serum from eight-week PBS or MDA-MB-231 EV treated mice was analyzed using CD32 and Angio 5-plex cytokine arrays (Eve Technologies, Calgary, Canada).

### Catalogue of Primary Antibodies and Dilutions

| Primary Antibodies | Cat # | Conc.<br>mg/ml | Dilution | Clone | Source |
| --- | --- | --- | --- | --- | --- |
| CD9 | 312102 | 0.5 | 1:1000 | HI9a | Biolegend |
| CD63 | cbl553 | 0.1 | 1:250 | RFAC4 | Millipore |
| CD63 | 353013 | 1 | 1:1000 | H5C6 | Biolegend |
| CD81 | 348501 | 0.5 | 1:1000 | 5A6 | Biolegend |
| ALIX | 634501 | 0.5 | 1:1000 | 3A9 | Biolegend |
| Flotillin-1 | 849801 | 0.5 | 1:1000 | W16108A | Biolegend |
| Calnexin | 699401 | 0.5 | 1:1000 | W17077 | Biolegend |
| Tsg101 | sc-136111 | 0.1 | 1:250 | 51 | Santa Cruz |
| MFGE8 | sc-271574 | 0.2 | 1:500 | F-5 | Santa Cruz |
| PE-mouse IgG1 k isotype | 400113 | 0.2 | 1:100 | MOPC-21 | Biolegend |
| PE anti-human CD63 | 353003 | 0.1 | 1:100 | H5C6 | Biolegend |
| PE anti-human NDPK-A | N7100-66A | 0.1 | 1:100 | - | US Biologicals |
| Rab anti-human NM23A | Ab92327 | 0.1 | 1:200 | EPR3036 | Abcam |
| Rab anti-human NME2 | Ab131329 | 0.1 | 1:200 | EPR8351 | Abcam |
| PE donkey anti-Rabbit | 406421 | 0.2 | 1:200 | Poly4064 | Biolegend |
| Rabbit IgG isotype | Ab172730 | 0.1 | 1:200 | EPR25A | Abcam |
| CPTC-NME1-5 | - | 0.03 | 1:1000 | - | DSHB |
| CPTC-NME2-1 | - | 0.47 | 1:1000 | - | DSHB |
| Firefly Luciferase | Ab21176 | 11.7 | 1:1000 | - | Abcam |
| Thrombospondin-1 | Ab85762 | 1 | 1:1000 | - | Abcam |
| GAPDH | D16H11 | - | 1:1000 | 5174 | CST |
| COX-1 | 4841S | - | 1:1000 | 9896 | CST |
| COX-2 | 4842S | - | 1:1000 | 12282 | CST |
| NOS2 (iNOS) | SC-650 | 0.2 | 1:1000 | M-19 | Santa Cruz |
| NOS3 (eNOS) | SC654 | 0.2 | 1:1000 | C-20 | Santa Cruz |
| ICAM-1 (CD54) | 116102 | 0.5 | 1:250 | YN1 | Biolegend |
| ICAM-2 (CD102) | 105601 | 0.5 | 1:250 | 3C4 | Biolegend |
| VCAM-1 (CD106) | 105702 | 0.5 | 1:250 | 429 | Biolegend |
| ZO-1 | D7D12 | - | 1:250 | 8193 | CST |
| β-catenin | D10A8 | - | 1:250 | 8480 | CST |
| Integrin β3 | SC-6627 | 0.2 | 1:1000 | N-20 | CST |
| GAPDH | SC-47724 | 0.2 | 1:1000 | 0411 | Santa Cruz |

## References

1. Siegel, R. L., Miller, K. D., Jemal, A. Cancer statistics, 2018. CA Cancer J Clin. 68, 7–30 (2018).

2. Kaplan, R.N. et al. VEGFR1-positive haematopoietic bone marrow progenitors initiate the pre-metastatic niche. Nature. 438, 820–827 (2005).

3. Hüsemann, Y. et al. Systemic spread is an early step in breast cancer. Cancer Cell. 13, 58–68 (2008).

4. Hosseini, H. et al. Early dissemination seeds metastasis in breast cancer. Nature. 10.1038/nature20785 (2016).

5. Harper, K. L et al. Mechanism of early dissemination and metastasis in Her2^+^ mammary cancer. Nature. 10.1038/nature20609 (2016).

6. Xu, R. et al. Extracellular vesicles in cancer — implications for future improvements in cancer care. Nat Rev Clin Oncology. 15, 617–638 (2018).

7. TKach, M., Thery, C. Communication by Extracellular Vesicles: Where We Are and Where We Need to Go. Cell. 164, 1226–1232 (2016).

8. Peinado, H. et al. Melanoma exosomes educate bone marrow progenitor cells toward a prometastatic phenotype through MET. Nature medicine. 18, 883–891 (2012).

9. Hoshino, A. et al. Tumour exosome integrins determine organotropic metastasis. Nature. 527, 329–335 (2015).

10. Steeg PS, et al. Evidence for a novel gene associated with low tumor metastatic potential. J Natl Cancer Inst. 1988;80(3):200–204.

11. Stahl JA, Leone A, Rosengard AM, et al. Identification of a second human nm23 gene, nm23-H2. Cancer Res. 1991;51:445–449.

12. Boissan M, Schlattner U, Lacombe ML. The NDPK/NME superfamily: state of the art. Lab Invest. 2018;98(2):164–174.

13. Sharma S, Sengupta A, Chowdhury S. NM23/NDPK proteins in transcription regulatory functions and chromatin modulation: emerging trends. Lab Invest. 2017;98(2):175–181.

14. Kruger, S. et al. Molecular characterization of exosome-like vesicles from breast cancer cells. BMC Cancer. 14, 44 (2014).

15. Palazzolo, G. et al. Proteomic analysis of exosome-like vesicles derived from breast cancer cells. Anticancer Res. 32, 847–860 (2012).

16. Hurwitz, S. N. et al. Proteomic profiling of NCI-60 extracellular vesicles uncovers common protein cargo and cancer type-specific biomarkers. Oncotarget. 7, 86999–87015 (2016).

17. Demory Beckler, M. et al. Proteomic Analysis of Exosomes from Mutant KRAS Colon Cancer Cells Identifies Intercellular Transfer of Mutant KRAS. Mol & Cell Proteomics: MCP. 12, 343–355 (2013).

18. Liang, B. et al. Characterization and proteomic analysis of ovarian cancer-derived exosomes. J Proteomics. 80, 171–182 (2013).

19. He, M. et al. Hepatocellular carcinoma-derived exosomes promote motility of immortalized hepatocyte through transfer of oncogenic proteins and RNAs. Carcinogenesis. 36, 1008–1018 (2015).

20. Lazar, I. et al. Proteome characterization of melanoma exosomes reveals a specific signature for metastatic cell lines. Pigment Cell Melanoma Res. 28, 464–75 (2015).

21. Buxton, I. L., Kaiser, R. A., Oxhorn, B. C., Cheek, D. J. Evidence supporting the Nucleotide Axis Hypothesis: ATP release and metabolism by coronary endothelium. Am J Physiol Heart Circ Physiol. 281, H1657–1666 (2001).

22. Di Virgilio, F., Sarti, A. C., Falzoni, S., De Marchi, E., Adinolfi, E. Extracellular ATP and P2 purinergic signalling in the tumour microenvironment. Nat Rev Canc. 18, 601–618 (2018).

23. Zhang, J. et al. ATP-P2Y2-β-catenin axis promotes cell invasion in breast cancer cells. Cancer Science. 108, 1318–1327 (2017).

24. Jin, H. et al. P2Y2 receptor activation by nucleotides released from highly metastatic breast cancer cells increases tumor growth and invasion via crosstalk with endothelial cells. Breast Cancer Res : BCR. 16, R77 (2014).

25. Adinolfi, E. et al. Expression of P2×7 receptor increases *in vivo* tumor growth. Cancer Res. 72, 2957–69 (2012).

26. Buvinic, S., Bravo-Zehnder, M., Boyer, J. L., Huidobro-Toro, J. P. Nucleotide P2Y1 receptor regulates EGF receptor mitogenic signaling and expression in epithelial cells. J Cell Sci. 120, 4289–4301 (2007).

27. Rumjahn, S. M., Javed, M. A., Wong, N., Law, W. E., Buxton, I. L. Purinergic regulation of angiogenesis by human breast carcinoma-secreted nucleoside diphosphate kinase. Br J Cancer. 97, 1372–1380 (2007).

28. Rumjahn, S. M., Yokdang, N., Baldwin, K. A., Thai, J., Buxton, I. L. Purinergic regulation of vascular endothelial growth factor signaling in angiogenesis. Br J Cancer. 100, 1465–1470 (2009).

29. Yokdang, N. et al. A role for nucleotides in support of breast cancer angiogenesis: heterologous receptor signalling. Br J Cancer. 104, 1628–1640 (2011).

30. Burnstock, G., & Ralevic, V. Purinergic signaling and blood vessels in health and disease. Pharmacol Rev. 66, 102–192 (2014).

31. Yokdang, N., Nordmeier, S., Speirs, K., Burkin, H. R., Buxton, I. L. Blockade of extracellular NM23 or its endothelial target slows breast cancer growth and metastasis. Integr Cancer Sci Ther. 2, 192–200 (2015).

32. Pellegatti, P. et al. Increased Level of Extracellular ATP at Tumor Sites: *In Vivo* Imaging with Plasma Membrane Luciferase. El Khoury J, ed. PLoS ONE. 3, e2599 (2008).

33. Joo, Y. N. et al. P2Y2R activation by nucleotides released from the highly metastatic breast cancer cell contributes to pre-metastatic niche formation by mediating lysyl oxidase secretion, collagen crosslinking, and monocyte recruitment. Oncotarget. 5, 9322–9334 (2014).

34. Sobczak, M., Dargatz, J., Chrzanowska-Wodnicka, M. Isolation and Culture of Pulmonary Endothelial Cells from Neonatal Mice. Journal of Visualized Experiments : JoVE. 2316 (2010).

35. Théry, C., Amigorena, S., Raposo, G., Clayton, A. Isolation and characterization of exosomes from cell culture supernatants and biological fluids. Curr Protoc Cell Biol Chapter. 3, Unit 3.22 (2006).

36. Martins-Green, M., Petreaca, M., Yao, M. An assay system for in vitro detection of permeability in human “endothelium”. Methods in enzymology. 443, 137–153 (2008).

37. Steeg, P. S. et al. Evidence for a novel gene associated with low tumor metastatic potential. J Natl Cancer Inst. 80, 200–204 (1988).

38. Otero-Estévez, O. et al. Evaluation of serum nucleoside diphosphate kinase A for the detection of colorectal cancer. Scientific Reports. 6, 26703 (2016).

39. Alvarez-Chaver, P. et al. Selection of putative colorectal cancer markers by applying PCA on the soluble proteome of tumors: NDK A as a promising candidate. J. Proteomics. 74, 874–886 (2011).

40. Okabe-Kado, J. et al. Clinical significance of serum NM23-H1 protein in neuroblastoma. Cancer Sci. 96, 653–660 (2005).

41. Su Kim, D. et al. Composite three-marker assay for early detection of kidney cancer. Cancer Epidemiol Biomarkers Prev. 22, 390–98 (2013).

42. Niitsu, N., Nakamine, H., Okamoto, M. Expression of nm23-H1 is associated with poor prognosis in peripheral T-cell lymphoma, not otherwise specified. Clin Cancer Res. 17, 2893–2899 (2011).

43. Yokdang, N., Buxton, N. D., Buxton, I. L. Measurement of human breast tumor cell-secreted shNDPK-B in a murine breast cancer model suggests its role in metastatic progression. Proc West Pharmacol Soc. 52, 88–91 (2009).

44. Anzinger, J., Malmquist, N. A., Gould, J., Buxton, I. L. Secretion of a nucleoside diphosphate kinase (Nm23-H2) by cells from human breast, colon, pancreas and lung tumors. Proc West Pharmacol Soc. 44, 61–63 (2001).

45. Ludwig, N., Yerneni, S. S., Razzo, B. M., Whiteside, T. L. Exosomes from HNSCC promote angiogenesis through reprogramming of endothelial cells. Mol Cancer Res. 16, 1798–1808 (2018).

46. Zhou, W. et al. Cancer-secreted miR-105 destroys vascular endothelial barriers to promote metastasis. Cancer Cell. 25, 501–515 (2014).

47. Hippe, H. J. et al. The interaction of nucleoside diphosphate kinase B with Gβγ dimers controls heterotrimeric G protein function. PNAS. 106, 16269–16274 (2009).

48. Feng, Y. et al. Nucleoside diphosphate kinase B regulates angiogenesis through modulation of vascular endothelial growth factor receptor type 2 and endothelial adherens junction proteins. Arterioscler Thromb Vasc Biol. 34, 2292–300 (2014).

49. Francois-Moutal, L. et al. Two-Step Membrane Binding of NDPK-B Induces Membrane Fluidity Decrease and Changes in Lipid Lateral Organization and Protein Cluster Formation. Langmuir ACS J Surf Colloids. 32, 12923–12933 (2016).

50. Gross, S. et al. Nucleoside diphosphate kinase B regulates angiogenic responses in the endothelium via caveolae formation and c-Src-mediated caveolin-1 phosphorylation. J Cereb Blood Flow Metab. 37, 2471–2484 (2016).

51. Jacobson, K. A., Deflorian, F., Mishra, S., Costanzi, S. Pharmacochemistry of the platelet purinergic receptors. Purinergic Signaling. 7, 305–324 (2011).

52. Labelle, M., Begum, S., Hynes, R. O. Platelets guide the formation of early metastatic niches. PNAS. 111, E3053–E3061 (2014).

53. Ward, Y. et al. Platelets Promote Metastasis via Binding Tumor CD97 Leading to Bidirectional Signaling that Coordinates Transendothelial Migration. Cell Reports. 23, 808–822 (2018).

54. Gomes, F. G. et al. Breast-cancer extracellular vesicles induce platelet activation and aggregation by tissue factor-independent and -dependent mechanisms. Thromb Res. 159, 24–32 (2017).

55. Blay J., White T. D., Hoskin D. W.. The extracellular fluid of solid carcinomas contains immunosuppressive concentrations of adenosine. Cancer Res. 57, 2602–5 (1997).

56. Pellegatti P., Raffaghello L., Bianchi G., Piccardi F., Pistoia V., Di V. F. Increased level of extracellular ATP at tumor sites: in vivo imaging with plasma membrane luciferase. PLoS One. 3, e2599 (2008).

57. de Andrade Mello P., Coutinho-Silva R., Savio L. E. B. Multifaceted Effects of Extracellular Adenosine Triphosphate and Adenosine in the Tumor-Host Interaction and Therapeutic Perspectives. Front Immunol. 8, 1526 (2017).

