## Supplementary material for "Extracellular Vesicle-Mediated Purinergic Signaling Contributes to Host Microenvironment Plasticity and Metastasis in Triple Negative Breast Cancer": All 8 supplemental figures and 4 supplemental tables

Supplemental Figure 1

Click here to access/download;Figure;Suplemental Fig 1.PNG

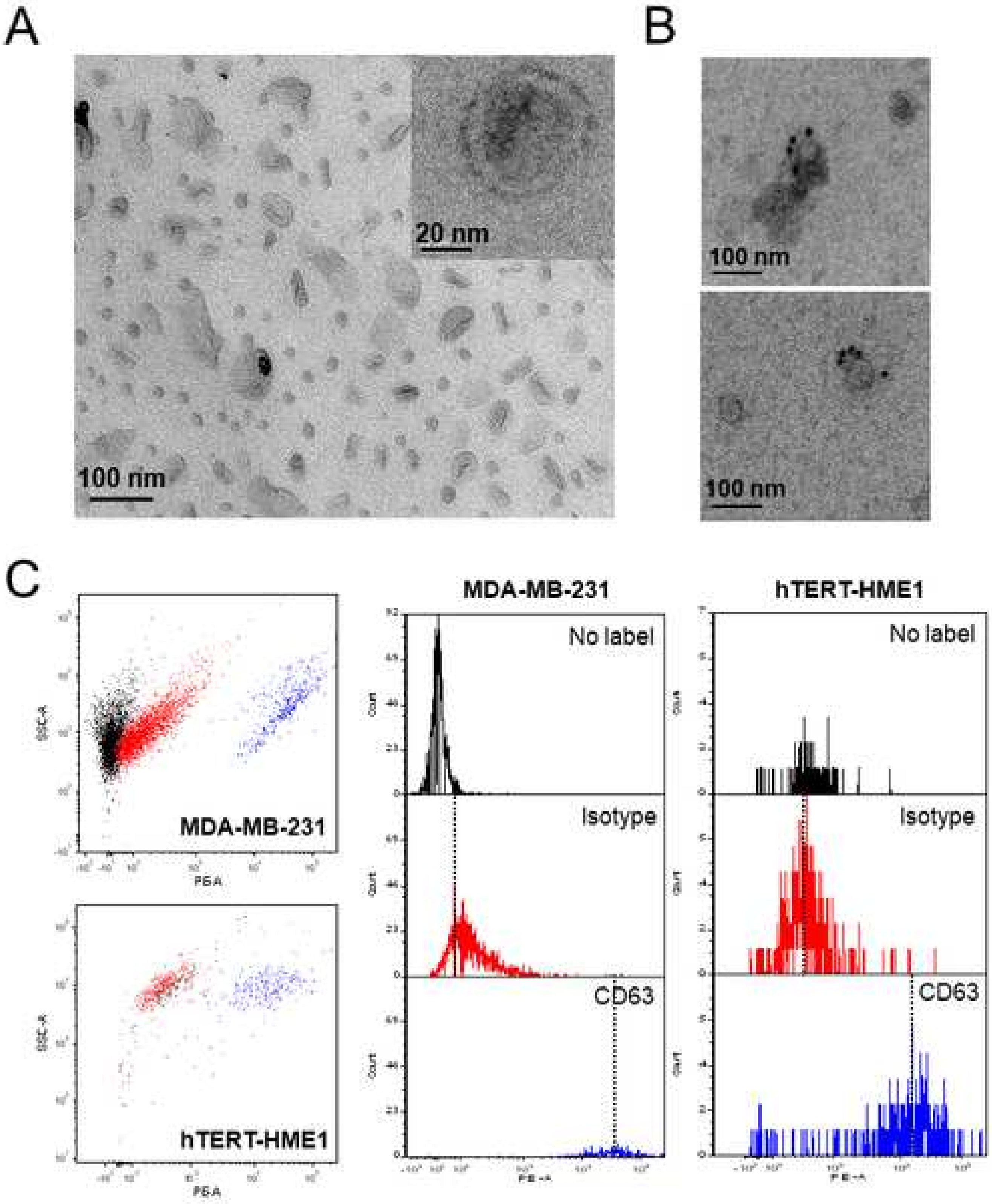


Supplemental 1

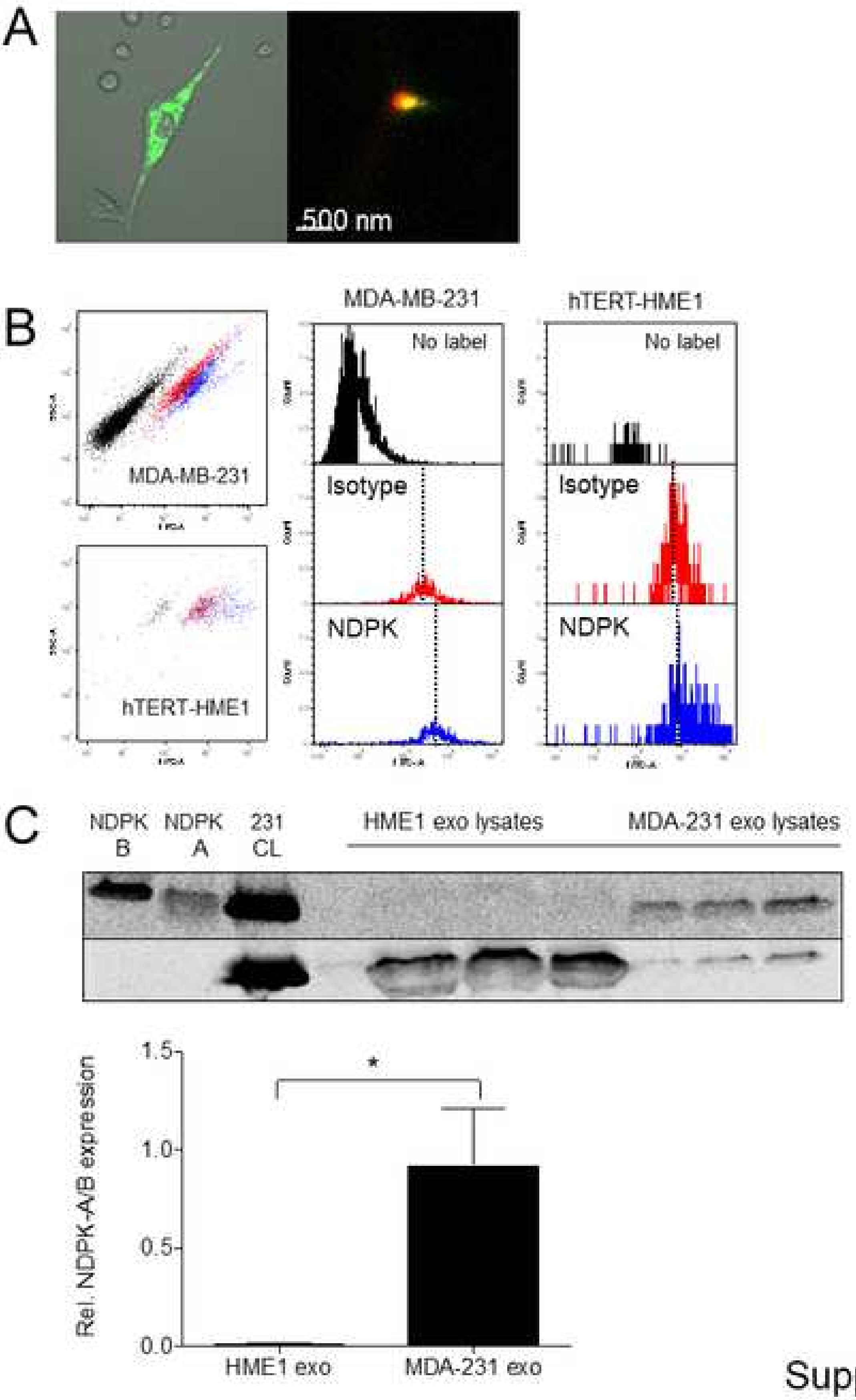


Supplemental 2a

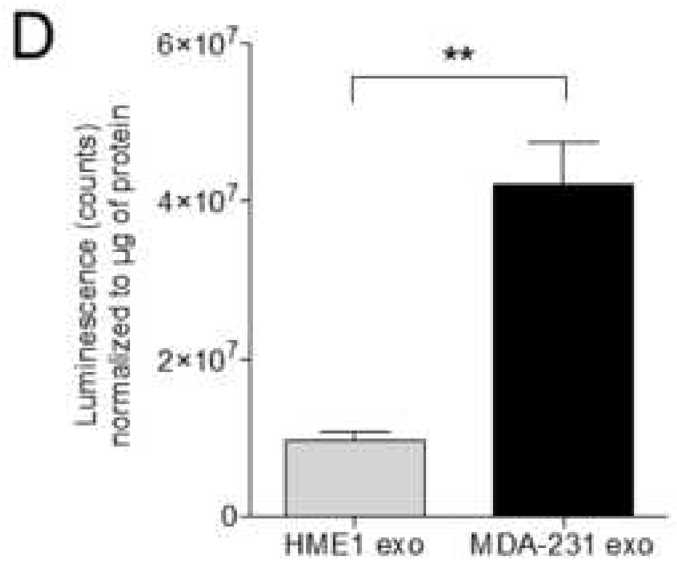


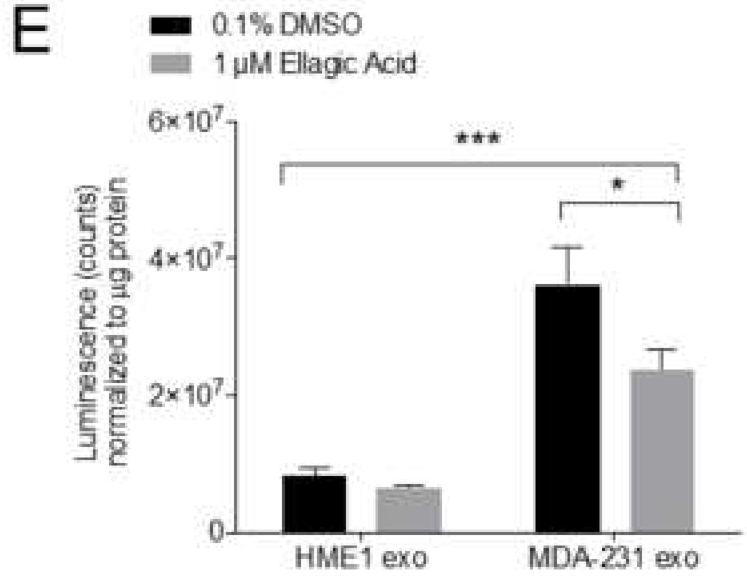


Supplemental 2b

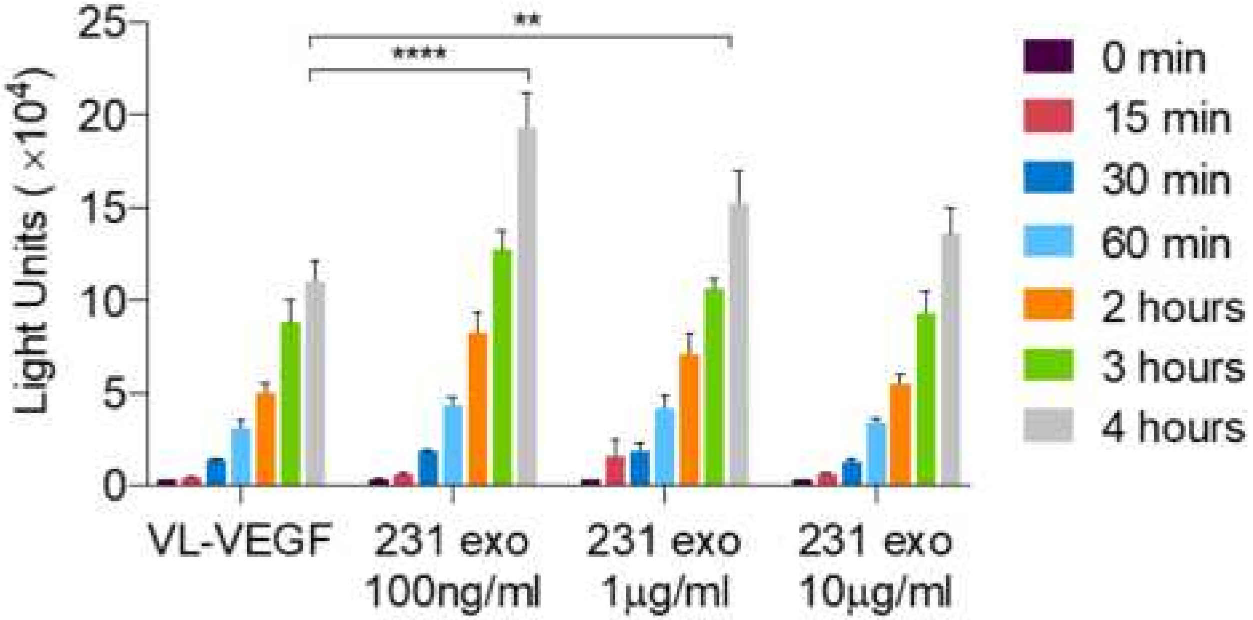


Supplemental 3

Supplemental Figure 4

Click here to access/download;Figure;Suplemental Fig 4.PNG

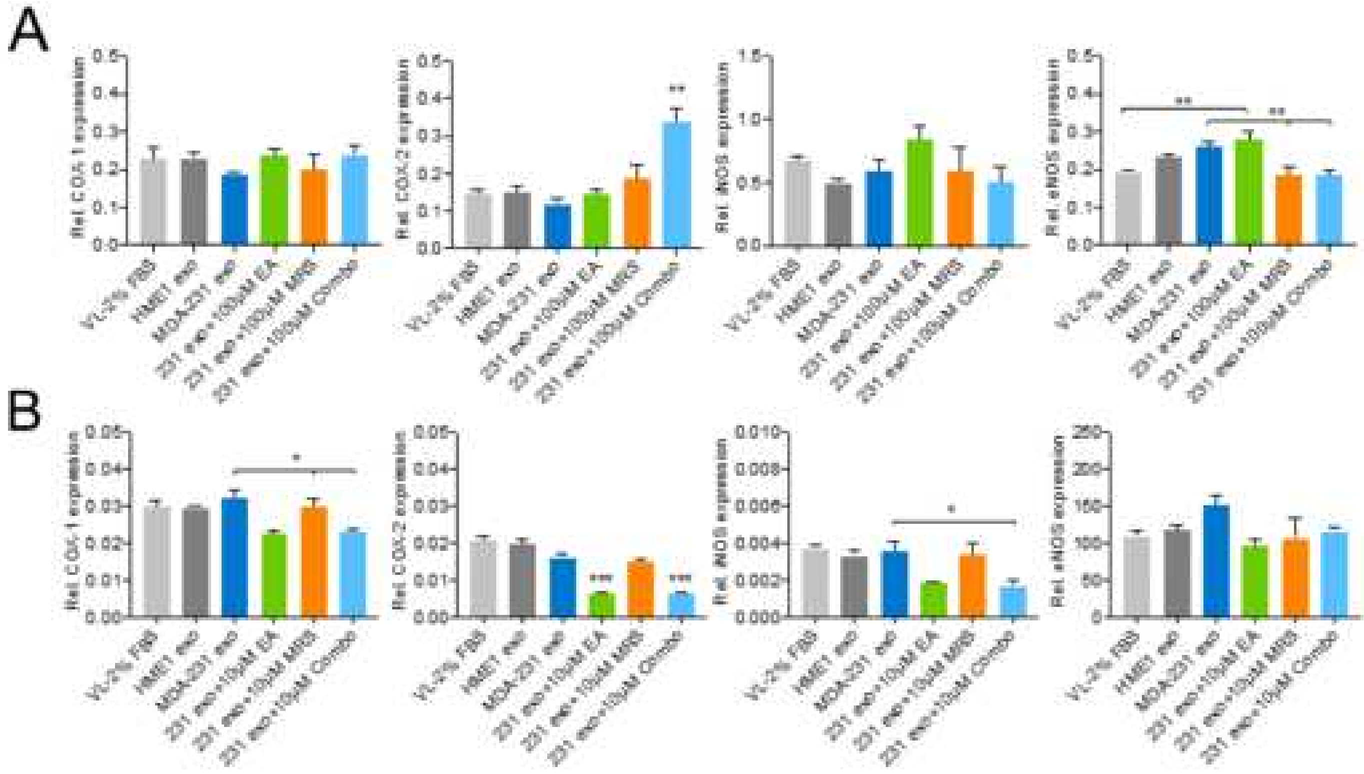

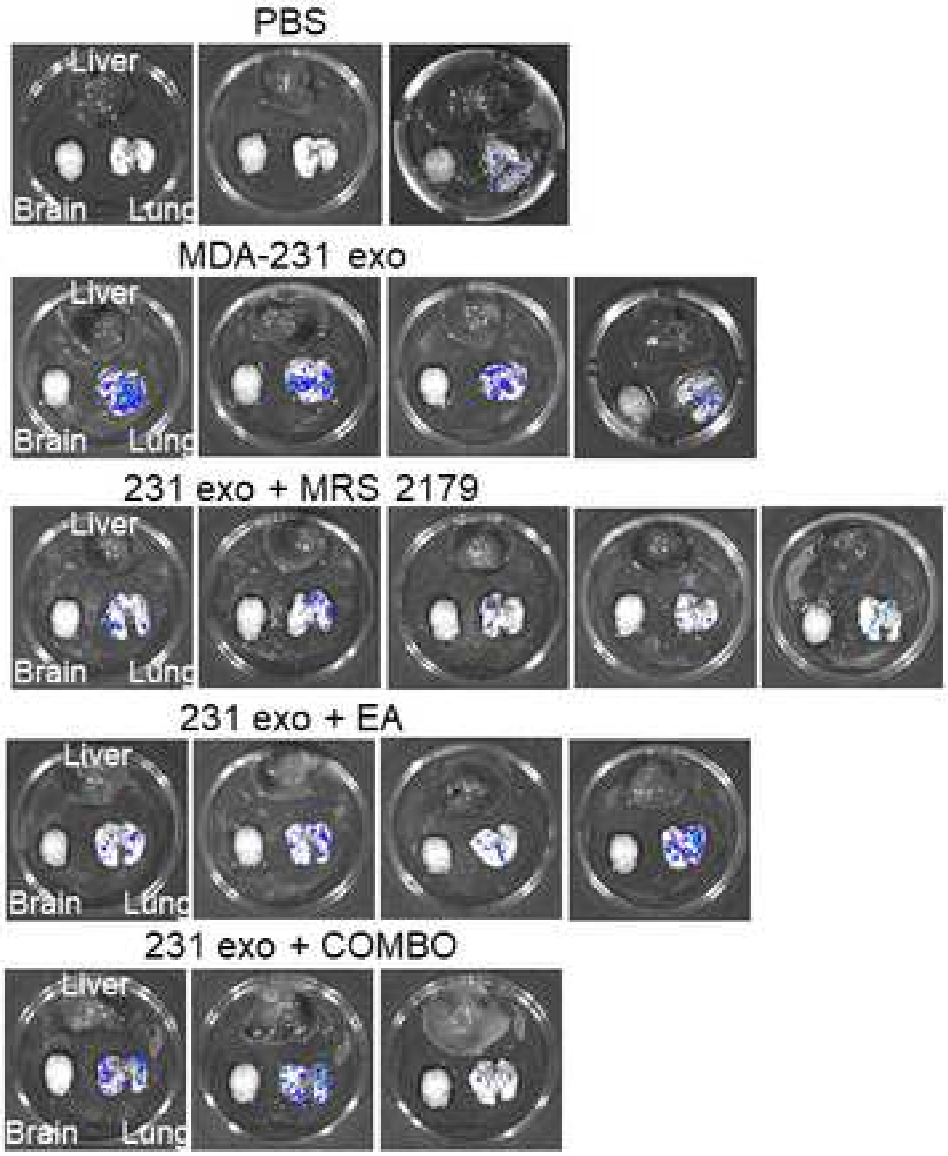
PBS
Liver
Brain
Lung
MDA-231 exo
231 exo + MRS 2179
231 exo + EA
231 exo + COMBO

Supplemental Figure 6
Click here to access/download;Figure;Suplemental Fig 6.PNG

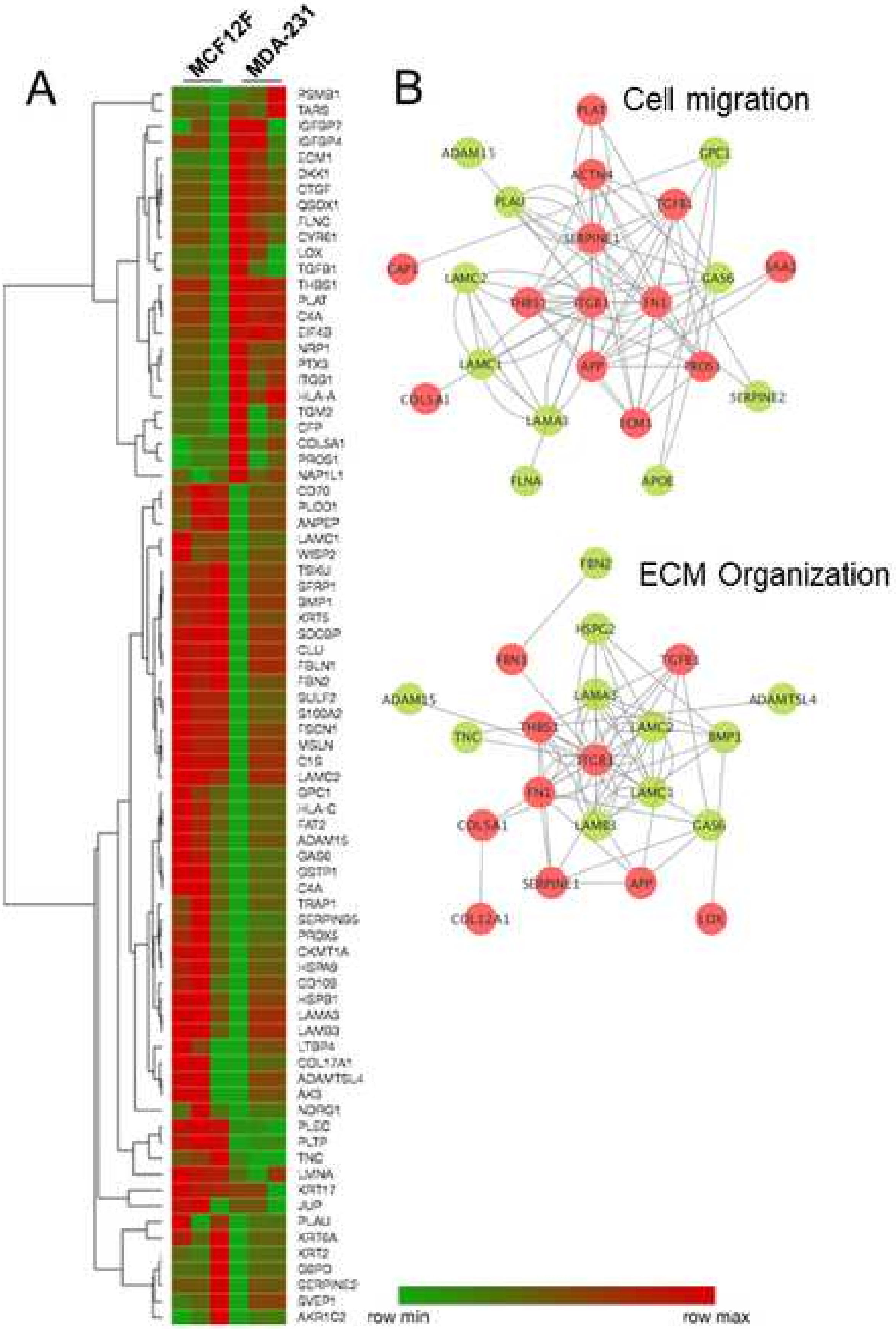


Supplemental 6

Click here to access/download;Figure;Suplemental Fig 7.PNG

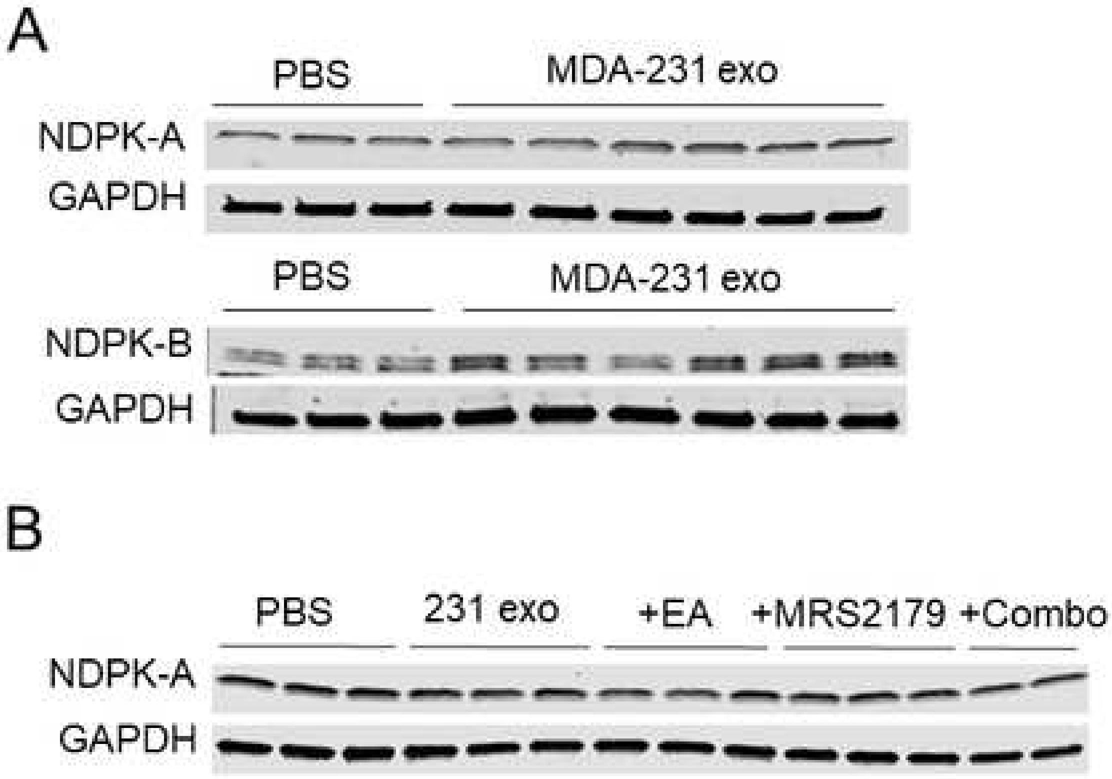


Supplemental 7

**Supplemental Table 1.** Proteins with greatest fold enrichment in MDA-231 exosomes

| Protein Name | Gene Name | log2 FC | P Value |
|---|---|---|---|
| Thrombospondin-1 | THBS1 | 7.1 | 9E-05 |
| Complement C4-A | C4A | 5.4 | 9E-05 |
| Insulin-like growth factor-binding protein 4 | IGFBP4 | 5.2 | 9E-05 |
| Protein CYR61 | CYR61 | 4.9 | 9E-05 |
| Tissue-type plasminogen activator | PLAT | 4.6 | 9E-05 |
| Connective tissue growth factor | CTGF | 4.2 | 9E-05 |
| Pentraxin-related protein PTX3 | PTX3 | 3.9 | 9E-05 |
| Sulfhydryl oxidase 1 | QSOX1 | 3.8 | 9E-05 |
| Transforming growth factor beta 1 | TGFB1 | 3.8 | 9E-05 |
| Neuropilin-1 | NRP1 | 3.8 | 9E-05 |
| Eukaryotic translation initiation factor 4B | EIF4B | 3.7 | 9E-05 |
| Integrin beta 1 | ITGB1 | 3.7 | 9E-05 |
| Filamin-C | FLNC | 3.7 | 9E-05 |
| Threonine-tRNA ligase | TARS | 3.6 | 9E-05 |
| Properdin | CFP | 3.5 | 9E-05 |
| Nucleosome assembly protein 1-like 1 | NAP1L1 | 3.5 | 3E-04 |
| Protein-lysine 6-oxidase | LOX | 3.3 | 1E-04 |
| Insulin-like growth factor-binding protein 7 | IGFBP7 | 3.3 | 9E-05 |
| Dickkopf-related protein 1 | DKK1 | 3.3 | 8E-04 |
| Protein-glutamine gamma-glutamyltransferase 2 | TGM2 | 3.2 | 3E-04 |

**Supplemental Table 2.** Proteins with greatest fold enrichment in MCF-12F exosomes

| Protein Name | Gene Name | log2 FC | P Value |
|---|---|---|---|
| Laminin subunit beta-3 | LAMB3 | 6.5 | 9E-05 |
| Laminin subunit alpha-3 | LAMA3 | 6.4 | 9E-05 |
| Laminin subunit gamma-2 | LAMC2 | 6.4 | 9E-05 |
| Fibulin-1 | FBLN1 | 6.3 | 9E-05 |
| Sushi, von Willebrand factor type A, EGF and pentraxin domain-containing protein1 | SVEP1 | 6.0 | 9E-05 |
| Clusterin | CLU | 5.6 | 9E-05 |
| Complement C1s subcomponent | C1S | 5.6 | 9E-05 |
| Mesothelin | MSLN | 5.2 | 9E-05 |
| Bone morphogenetic protein 1 | BMP1 | 5.1 | 9E-05 |
| Latent-transforming growth factor beta-binding protein 1 | LTBP4 | 4.8 | 9E-05 |
| Syntenin-1 | SDCBP | 4.8 | 9E-05 |
| ADATS-like protein 4 | ADAMTSL4 | 4.8 | 9E-05 |
| Secreted frizzled-related protein 1 | SFRP1 | 4.7 | 9E-05 |
| Keratin, type II cytoskeletal 17 | KRT17 | 4.6 | 9E-05 |
| Aminopeptidase | ANPEP | 4.6 | 9E-05 |
| Procollagen-lysine, 2-oxoglutarate 5-dioxygenase 1 | PLOD1 | 4.6 | 9E-05 |
| Heat shock protein beta-1 | HSPB1 | 4.6 | 9E-05 |
| Glia-derived nexin | SERPINE2 | 4.5 | 9E-05 |
| CD109 antigen | CD109 | 4.3 | 9E-05 |
| Tsukushin | TSKU | 4.2 | 5E-04 |

**Supplemental Table 3.** Top up-regulated lung proteins after MDA-MB-231 exosome treatment

| Protein Name | Gene Name | log2 FC | P Value |
|---|---|---|---|
| Alpha-1-antitrypsin 1-5 | Serpina1e | 1.00 | 0.0404 |
| Interferon-inducible GTPase 1 | Iigp1 | 0.89 | 0.0110 |
| Glycoprotein Ib, beta polypeptide | Gp1bb | 0.83 | 0.0388 |
| Solute carrier family 2, facilitated glucose transporter member 3 | Slc2a3 | 0.77 | 0.0164 |
| Interferon-induced protein 44-like | Ifi44l | 0.77 | 0.0021 |
| FYN-binding protein | Fyb | 0.76 | 0.0228 |
| Tubulin beta-1 chain | Tubb1 | 0.72 | 0.0091 |
| Integrin alpha-Iib | Itga2b | 0.72 | 0.0083 |
| Integrin beta-3 | Itgb3 | 0.71 | 0.0041 |
| C-type lectin domain family 1 member B | Clec1b | 0.70 | 0.0035 |
| Lymphocyte cytosolic protein 2 | Lcp2 | 0.69 | 0.0027 |
| Thrombospondin 1 | Thbs1 | 0.68 | 0.0048 |
| Bridging integrator 2 | Bin2 | 0.66 | 0.0067 |
| Metalloreductase STEAP4 | Steap4 | 0.64 | 0.0028 |
| Platelet glycoprotein IX | Gp9 | 0.61 | 0.0344 |
| Fermitin family homolog 3 | Fermt3 | 0.58 | 0.0029 |
| Phospholipase D4 | Pld4 | 0.58 | 0.0003 |
| DNA replication licensing factor MCM3 | Mcm3 | 0.57 | 0.0226 |
| Multimerin-1 | Mmrn1 | 0.56 | 0.0361 |
| DNA replication licensing factor MCM4 | Mcm4 | 0.55 | 0.0238 |

**Supplemental Table 4.** Top down-regulated lung proteins after MDA-231 exosome treatment

| Protein Name | Gene Name | log2 FC | P Value |
|---|---|---|---|
| Migration and invasion enhancer 1 | Mien1 | -0.34 | 0.0047 |
| Stonin-1 | Ston1 | -0.37 | 0.0254 |
| Polymeric immunoglobulin receptor | Pigr | -0.32 | 0.0145 |
| PTB domain-containing engulfment adapter protein 1 | Gulp1 | -0.29 | 0.0006 |
| Voltage-dependent calcium channel subunit alpha-2/delta-1 | Cacna2d1 | -0.29 | 0.0139 |
| Carboxylic ester hydrolase | Ces2g | -0.26 | 0.0021 |
| UPF0585 protein C16orf13 homolog | | -0.25 | 0.0193 |
| CD82 antigen | Cd82 | -0.25 | 0.0408 |
| Hepatocyte growth factor activator | Hgfac | -0.24 | 0.0357 |
| Alkaline phosphatase, tissue-nonspecific isozyme | Alpl | -0.24 | 0.0573 |
| Tumor-associated calcium signal transducer 2 | Tacstd2 | -0.23 | 0.0040 |
| CDP-diacylglycerol--inositol 3-phosphatidyltransferase | Cdipt | -0.23 | 0.0085 |
| Annexin A9 | Anxa9 | -0.23 | 0.0225 |
| Angiotensin-converting enzyme | Ace | -0.23 | 0.0595 |
| Septin-5 | Sept5 | -0.22 | 0.0164 |
| Heat shock protein beta-8 | Hspb8 | -0.22 | 0.0626 |
| Transforming growth factor beta receptor type 3 | Tgfbr3 | -0.22 | 0.0008 |
| Serine/threonine-protein phosphatase 2A | Ppp2r5c | -0.22 | 0.0259 |
| Epoxide hydrolase 1 | Ephx1 | -0.21 | 0.0046 |
| Claudin-5 | Cldn5 | -0.21 | 0.0031 |
